# Poly-ADP-ribosylation drives loss of protein homeostasis in ATM and Mre11 deficiency

**DOI:** 10.1101/2020.10.27.357210

**Authors:** Ji-Hoon Lee, Seung W. Ryu, Nicolette A. Ender, Tanya T. Paull

## Abstract

Loss of the ataxia-telangiectasia mutated (ATM) kinase causes cerebellum-specific neurodegeneration in humans. We previously demonstrated that deficiency in ATM activation via oxidative stress generates high levels of insoluble protein aggregates in human cells, reminiscent of protein dysfunction in common neurodegenerative disorders. Here we show that this process is driven by poly-ADP-ribose polymerases (PARPs) and that the insoluble protein species arise from intrinsically disordered proteins associating with PAR-associated genomic sites in ATM-deficient cells. The lesions implicated in this process are single-strand DNA breaks dependent on reactive oxygen species, transcription, and R-loops. Human cells expressing Mre11 A-T-like disorder (ATLD) mutants also show PARP-dependent aggregation identical to that of ATM deficiency. Lastly, analysis of A-T patient cerebellum samples shows widespread protein aggregation as well as loss of proteins known to be critical in human spinocerebellar ataxias. These results provide a new hypothesis for loss of protein integrity and cerebellum function in A-T.

## Introduction

Ataxia-telangiectasia Mutated (ATM) is a master regulator of the DNA damage response in eukaryotes, responding to DNA double-strand breaks within seconds and phosphorylating several hundred targets to promote checkpoint activation, DNA repair, and many other processes related to DNA metabolism (Shiloh and Ziv, 2013). Loss of ATM in humans causes the autosomal recessive disorder ataxia-telangiectasia (A-T), a pleiotropic disease that includes a high frequency of malignancy and immunodeficiency. These features are consistent with the important role of ATM in regulation of the cell cycle and repair in response to DNA damage, particularly during immunoglobulin gene rearrangements in the immune system (Cremona and Behrens, 2014; Ghosh et al., 2018).

The primary clinical feature of A-T patients, however, is the early-onset cerebellar neurodegeneration that appears in the first few years of life and progressively worsens through early adulthood (Rothblum-Oviatt et al., 2016). The ataxia that results from this cerebellum-specific neurodegeneration in A-T patients has some similar features to that seen in other familial ataxias, including spinocerebellar ataxias (SCAs), a heterogeneous group of mostly dominantly inherited syndromes (Klockgether et al., 2019). A subset of SCAs result from trinucleotide expansion mutations in translated regions of genes, and several SCA mutant alleles have been shown to express proteins that form insoluble aggregates in mammalian cells, a property that is linked to their pathology (Seidel et al., 2012).

The source of neurotoxicity in A-T has been widely debated and is still unclear. Previous work in the field has highlighted the potential role of oxidative stress in A-T, showing that cells from A-T patients have high levels of reactive oxygen species (ROS) and signs of chronic oxidative stress, that ROS-mediated signaling in ATM-deficient cells is aberrant, and that mouse models of ATM deficiency lose stem cell populations due to insufficient antioxidant capacity (Barlow et al., 1999; Barzilai et al., 2002; Ditch and Paull, 2011; Ito et al., 2004). Several years ago we demonstrated that ATM can be directly activated by oxidative stress, independent of DNA damage, by the formation of disulfide bonds between the two monomers in an ATM dimer (Guo et al., 2010). We identified mutations that specifically block this oxidative stress pathway, one of which is associated with neurodegeneration in several A-T patients, R3047X (Chessa et al., 1992; Gilad et al., 1998; Toyoshima et al., 1998). The cerebellar degeneration observed in this subset of A-T “variants” suggests that loss of the oxidative activation pathway generates the neurodegenerative aspects of the phenotype.

Another subset of patients, those with A-T-like disorder (ATLD), are difficult to reconcile with this hypothesis, however. ATLD is caused by mutations in the gene encoding Mre11, a component of the Mre11-Rad50-Nbs1 (MRN) complex that plays a key role in activating ATM in the DNA damage response (Lee and Paull, 2005; Stewart et al., 1999; Stracker and Petrini, 2011). Rare hypomorphic mutations in Mre11 generate a clinical phenotype strongly overlapping with that of A-T, including cerebellar neurodegeneration (but no telangiectasia and varying cancer incidence)(Regal et al., 2013; Taylor et al., 2004). ATLD mutations were originally shown to compromise ATM activation by double-strand breaks, a result generally interpreted to mean that the ataxia observed in A-T and ATLD must be due to loss of MRN activation of ATM. This is also an attractive model but is difficult to reconcile with the oxidative stress hypothesis discussed above.

To investigate ATM-dependent signaling pathways with greater specificity we previously identified alleles of ATM deficient in MRN-dependent activation by double-strand breaks, in addition to the ROS-specific separation of function alleles (Lee et al., 2018). We generated an inducible expression system for recombinant ATM in human cells combined with shRNA depletion of the endogenous protein and showed that the MRN-dependent pathway is essential for survival of DNA damage, DNA end processing, and damage-induced checkpoints and autophagy, while the oxidation-dependent pathway is essential for regulating ROS levels in human cells as well as ROS-induced autophagy and mitophagy. We also found that cells expressing ATM deficient in ROS activation, as well as A-T patient cells, exhibit widespread protein aggregation that is exacerbated by additional oxidative stress.

We focused on this apparent dysregulation of protein homeostasis in A-T cells because many of the more common neurodegenerative diseases in the human population are associated with aggregation of specific polypeptides in the brain, in some cases with clear causative links between the aggregation and neurotoxicity, and diverse model systems have confirmed these relationships (Bourdenx et al., 2017; Currais et al., 2017; Gidalevitz et al., 2006; Groh et al., 2017; Kikis et al., 2010; Ross and Poirier, 2004). In addition, several of the SCA ataxias, while associated with diverse genetic mutations, generate aggregation-prone mutant proteins that cause cerebellum-specific neurodegeneration similar to that observed in A-T (Buijsen et al., 2019; Klockgether et al., 2019).

In the work presented here we investigate the source of the protein aggregation observed in the absence of ATM function and find that it is dependent on poly(ADP-ribose) polymerase (PARP) activation at sites of single-strand breaks in genomic DNA. The breaks are transcription-dependent and exacerbated by elevated ROS generated when ATM oxidative activation is blocked or when ATM is absent. We find that PARP-dependent aggregates also appear in ATLD cells, indicating that loss of Mre11 function and ATM deficiency have similar endpoints with respect to protein homeostasis. Finally, we show that widespread protein aggregation is also clearly apparent in A-T patient cerebellum tissue compared to control samples, and is correlated with the loss of several proteins implicated in familial SCA. These findings lead us to a new view of the connection between DNA damage and pathological self-assembly of intrinsically disordered proteins, and suggest a model for understanding the progression of events that drives cerebellum-specific neurodegeneration in A-T patients.

## Results

### Destabilization of proteins in human cells induced by ATM deficiency

We previously demonstrated widespread protein aggregation in the absence of ATM function in several different human cell lines, including A-T patient lymphoblasts, HEK293 embryonic kidney cells, and U2OS osteosarcoma cells that was apparent in the absence of treatment and accentuated with additional oxidative stress (Lee et al., 2018). Here we used U2OS cells to identify proteins that aggregate in the absence of ATM function as we did previously, comparing changes in the total proteome (lysate) to proteins identified in the detergent-resistant fraction (aggregates)(Fig. 1). Isolation of the aggregates involves multiple rounds of intensive sonication, solubilization, and centrifugation, resulting in a final detergent-resistant aggregate fraction. Of 2,363 proteins identified by label-free mass spectrometry in the total lysate, none showed significant differences in control versus ATM-depleted cells (Fig. 1A, Table S1), both treated with a low dose of arsenite. In the aggregate fraction, however, 1584 of these proteins showed differences in ATM-depleted cells, after controlling for false discovery rate with the Benjamini-Hochberg method (Fig. 1B, Table S1). Of these, the vast majority (1,579) show higher levels of aggregation in ATM-depleted cells. Among this group of destabilized proteins is PSMB2, a subunit of the proteasome, and CK2β, a subunit of the CK2 kinase that we monitored extensively in previous work (Lee et al., 2018).

**Figure 1.**
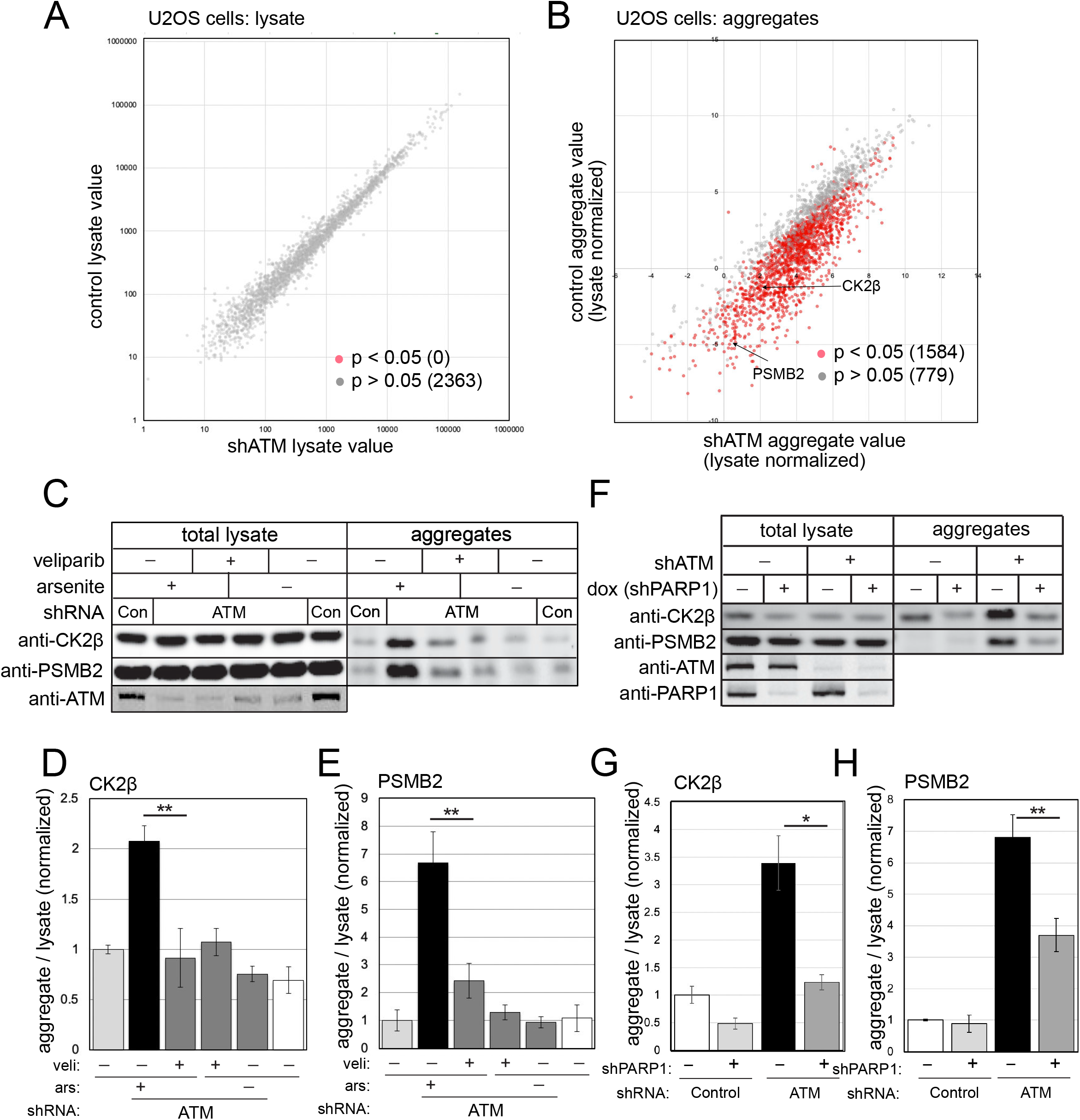
PARylation promotes protein aggregation in human cells lacking ATM. (A) Human U2OS cells were grown in the presence of arsenite (25 μM) and were treated with either control conditions or depleted of ATM by shRNA (3 biological replicates for each). Cell lysates were prepared and analyzed by mass spectrometry, identifying 2363 polypeptides in all samples. The levels of each protein in control versus ATM-depleted cells are shown, with non-significant differences in grey (2363) and significant differences in red (0), after FRD control at 0.05. (B) Detergent-resistant aggregates were prepared from control and ATM-depleted cells described in (A), and characterized by mass spectrometry. The levels of each protein in control versus ATM-depleted cells are shown, with non-significant differences in grey (779) and significant differences in red (1564), after FDR control at 0.05. (C) U2OS cells with control or ATM shRNA treatment were grown in the presence of arsenite (25 μM) and 10 μM veliparib as indicated. Lysates were prepared after 1 day of treatment as well as detergent-resistant aggregate fractions, which were analyzed by western blotting for CK2β or PSMB2. (D, E) Three replicates of the experiment shown in (C) were performed and quantified. Levels of CK2β (D) or PSMB2 (E) in aggregate fractions normalized by lysate levels were quantified from each experiment and shown here relative to control cells. (F) U2OS cells were depleted of ATM by shRNA and treated with 25 μM arsenite and shRNA directed against PARP1, as indicated. PARP1 shRNA expression is inducible with doxycycline. Levels of ATM, PARP1, CK2β and PSMB2 in lysates and aggregate fractions were analyzed by western blotting. (G, H) Three replicates of the experiment shown in (F) were performed and quantified. Levels of CK2β (G) or PSMB2 (H) in aggregate fractions normalized by lysate levels were quantified from each experiment and shown here relative to control cells. *, **, ***, and **** indicate p<0.05, 0.005, and 0.0005 by student t test; NS = not significant.

In addition to U2OS cells, we also examined protein aggregation in a human brain-derived glioblastoma cell line, U87-MG. Here we also depleted ATM with shRNA and found that aggregates were apparent in the absence of any exogenous stress (Fig. S1, Table S2). Mass spectrometry analysis of the total lysates did not detect any significant differences with ATM depletion, but we identified 560 proteins showing aggregation with ATM depletion, of which all showed higher levels in the depleted cells compared to the control. As in U2OS cells (Lee et al., 2018), these aggregates were completely eliminated with anti-oxidant treatment (N-acetyl cysteine, NAC)(Fig. S1). 84% of the aggregates identified in U87-MG cells also were identified as ATM-sensitive in U2OS, indicating that, despite the differences in cell origin, there is a reproducible pattern of polypeptides destabilized with ATM depletion.

### Protein aggregation observed with ATM depletion is dependent on PARP activity

PARylation of proteins by poly-ADP-ribose polymerases (PARPs) at sites of DNA single-strand and double-strand breaks is known to generate clouds of negatively-charged PAR polymers which have been shown by other groups to attract intrinsically disordered proteins (Altmeyer et al., 2015; Kai, 2016). PARylation has also been suggested to increase in ATM-deficient cells (Fang et al., 2016). Based on these observations, we reasoned that PARP may be involved in the aggregation of disordered proteins in ATM-deficient cells. To test this hypothesis, we used the PARP inhibitor veliparib, specific for the dominant PARP enzyme PARP1, as well as PARP2 (Knezevic et al., 2016). Here we found that incubation of U2OS cells with veliparib reduces the CK2β and PSMB2 aggregates observed with ATM depletion and arsenite treatment (Fig. 1C, quantification in Fig. 1D, E). To validate this result, we also depleted PARP1 with shRNA and found that this reduced aggregate species substantially (Fig. 1F, quantified in 1G, H). PARP2 depletion reduces aggregates as well, although to a lower extent than PARP1 depletion (Fig. S2; specificity of depletions for PARP1 and 2 also shown). Taken together, this evidence suggests that protein aggregation that we have monitored in ATM-depleted cells is PARP-dependent.

To quantify differences in PARylation in ATM-depleted cells, we utilized a recently developed live-cell PARylation sensor (Krastev et al., 2018)(Fig. 2) that fuses PAR-binding domains (PBZ) to split Venus proteins. In this system, simultaneous association of the PBZ-Venus(N-term) and PBZ-Venus(C-term) proteins at sites of PARylation is required for productive formation of the green fluorescent Venus protein. We expressed the PARylation sensor in U2OS cells, where it generates a nuclear punctate signal (Fig. 2A). To remove any non-stably associated Venus protein, we used detergent extraction of cells (which also removes the cytoplasm). The fluorescence yield per cell was measured by fluorescence-activated cell sorting (FACS), with approximately 10,000 cells monitored per measurement (3 biological replicates shown here). As shown in Fig. 2B, incubation of cells with arsenite alone increased the level of PAR sensor fluorescence, which increased further with concurrent ATM inhibition (AZD1390)(Durant et al., 2018). The level of PARylation observed with ATM depletion was reduced by veliparib treatment, as expected, since the majority of PAR formed in cells is generated by PARP1 and PARP2 (Gupte et al., 2017). PARylation as measured with this sensor was also reduced by NAC treatment in U2OS cells (Fig. 2C). In U87-MG cells, we also observed that the level of PARylation was reduced by both veliparib and NAC (Fig. S1).

**Figure 2.**
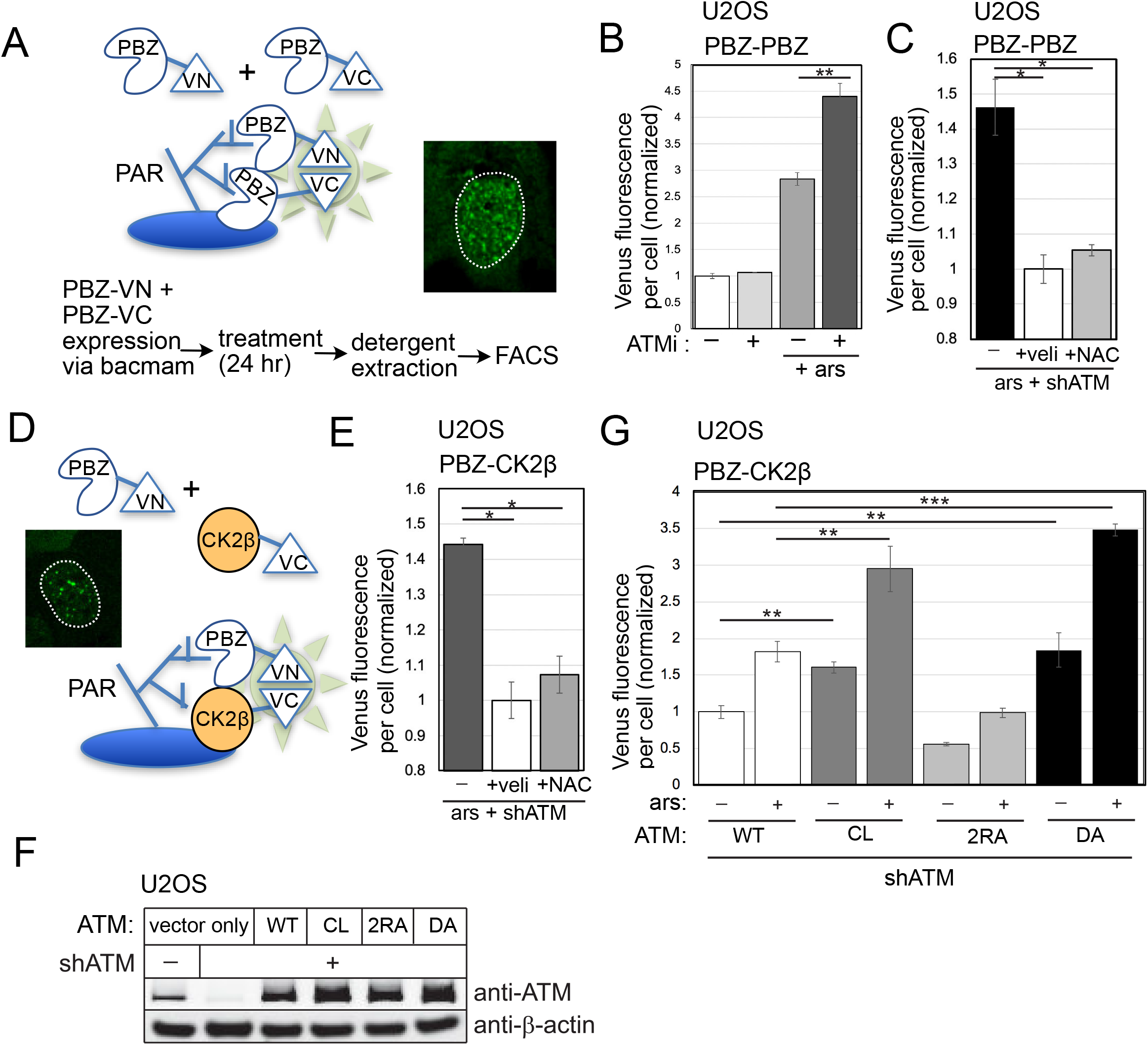
Live-cell sensors for PARylation show higher levels with ATM depletion. (A) Diagram of PBZ-PBZ live-cell split Venus sensor, adapted from (Krastev et al., 2018), with summary of workflow and inset of fluorescence signal from U2OS cells with ATM inhibitor treatment. (B) FACS results from three replicates showing the mean fluorescence yield per cell from cells expressing the PBZ-PBZ PAR sensor with ATM inhibitor (ATMi, 1 μM AZD1390) and arsenite (25 μM) as indicated. At least 10,000 cells were measured in each replicate. Fluorescence yield was normalized to control cells. (C) FACS results from three replicates showing the mean fluorescence yield per cell from cells expressing the PBZ-PBZ PAR sensor with arsenite, ATM shRNA, veliparib (10 μM), and NAC (1 mM) treatment as indicated. Fluorescence yield was normalized to veliparib-treated cells. (D) Diagram of PBZ-CK2β live-cell split Venus sensor, similar to the PBZ-PBZ sensor but with CK2β replacing the PBZ domain fused to VC, with inset of fluorescence signal from U2OS cells with ATM depletion. (E) FACS results from three replicates showing the mean fluorescence yield per cell from cells expressing the PBZ-CK2β sensor with arsenite, ATM shRNA, veliparib, and NAC treatment as indicated. Fluorescence yield was normalized to veliparib-treated cells. (F) FACS results from three replicates showing the mean fluorescence yield per cell from cells expressing the PBZ-CK2β sensor in U2OS cells with depletion of endogenous ATM and expression of recombinant WT, CL, 2RA, or DA alleles, with arsenite as indicated. At least 10,000 cells were measured in each replicate. Fluorescence yield was normalized to cells expressing WT ATM. Error bars indicate standard deviation. *, **, ***, and **** indicate p<0.05, 0.005, and 0.0005 by student t test; NS = not significant.

A subset of PARylation in actively growing cells derives from nicks associated with Okazaki fragments (Hanzlikova et al., 2018). Expression of the mutant or wild-type ATM alleles does not significantly alter the cell cycle in U2OS cells (data not shown), but we also asked whether the effect of ATM loss on PAR accumulation also occurs in non-dividing cells. To do this, we incubated U2OS cells in serum-free media, allowing them to reach a non-dividing state after 7 days. Measurement of the PBZ-PBZ sensor under these conditions showed significantly lower levels of PARylation overall, but still revealed elevated levels of fluorescence with ATM inhibition similar to actively dividing cells (Fig. S3).

We reasoned that the intrinsically disordered and oxidized proteins forming the aggregates could be assembling at the sites of PARP1/2-mediated PARylation. We tested this hypothesis directly by modifying the PAR sensor to include a disorder-prone protein in place of one of the PBZ domains (see Fig. 2D). We used the oxidation-prone CK2β protein for this purpose since it has been a reliable marker for aggregation in our experiments, and fused this protein to the C-terminal domain of the split Venus fluorescent protein. Using this hybrid PBZ-CK2β sensor, we found that ATM inhibition increased the level of PAR-CK2β association, and that NAC reduces the fluorescence yield significantly (Fig. 2E).

We then tested whether the ATM separation-of-function mutants we have previously described affect the association between CK2β and PAR. In this experiment, endogenous ATM was depleted and the WT, CL, 2RA, and kinase-dead D2889A (DA) (Daniel et al., 2012) alleles of ATM were inducibly expressed (Fig. 2F). The C2991L (CL) mutant of ATM is deficient in ROS-mediated activation (Guo et al., 2010; Lee et al., 2018) while the 2RA allele is deficient in MRN-mediated activation (Lee et al., 2018). The PBZ-CK2β sensor showed significantly higher levels of fluorescence, both with and without arsenite, with expression of the CL allele compared to WT ATM (Fig. 2G). In contrast, the 2RA mutant form of ATM showed levels of PBZ-CK2β Venus fluorescence even lower than in cells with WT ATM expressed, while the kinase-deficient DA allele was similar to the CL mutant.

### Transcription-dependent lesions are responsible for protein aggregation in ATM-deficient cells

The hyperactivation of PARP observed in ATM-deficient cells suggests the presence of DNA lesions. We have not observed higher levels of spontaneous double-strand breaks in the absence of ATM function (data not shown), but previous work has suggested that DNA lesions other than double-strand breaks may accumulate in the absence of ATM function (Katyal et al., 2014; Yamamoto et al., 2016). We examined levels of total single-strand DNA breaks using an alkaline comet assay in human U2OS cells depleted of endogenous ATM with concurrent expression of WT, CL, or R3047X (RX) alleles of ATM (Fig. 3A). The R3047X mutant form of ATM is deficient in ROS activation in vitro and in human cells, similar to the CL allele (Guo et al., 2010; Lee et al., 2018), and is also an A-T patient allele (Chessa et al., 1992; Gilad et al., 1998; Uhrhammer et al., 2002). We found that expression of either allele increased the levels of single-strand breaks significantly in comparison to the WT enzyme (Fig. 3B, C). This pattern is consistent with the increased ROS observed with both of these alleles, measured here using the oxidation-sensitive fluorogenic probe CellROX Deep Red (Fig. 3D). We found that addition of the antioxidant N-acetyl cysteine (NAC) eliminated the increased strand breaks observed with the mutant ATM (Fig. 3C), indicating that ROS is an important factor driving the appearance of the lesions.

**Figure 3.**
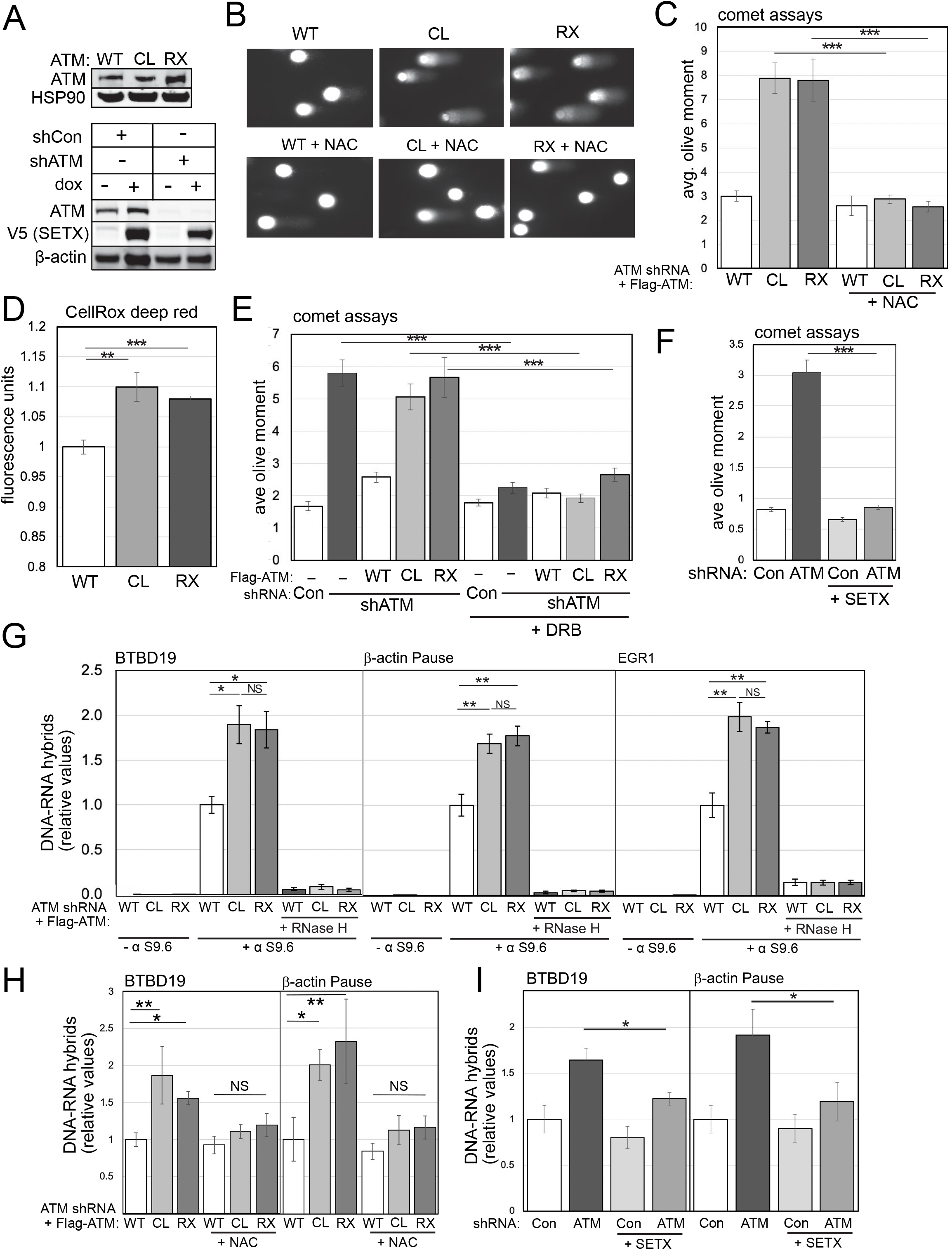
ATM-deficient cells accumulate single-strand breaks and R-loops. (A) Upper panel: Human U2OS cells were depleted of ATM with shRNA and wild-type (WT), C2991L (CL), or R3047X (RX) ATM alleles, shown by western blotting with anti-ATM antibody, with HSP90 as a blot control. Lower panel: U2OS cells were depleted of ATM with shRNA and the V5-tagged C-terminus of Senataxin was inducibly expressed with doxycycline, as indicated. Western blot shows levels of ATM and SETX (with V5 antibody), with β-actin as a loading control. (B) Alkaline comet assays were performed in U2OS cells depleted of endogenous ATM and expressing WT, CL, or RX alleles of ATM with NAC treatment (1mM) as indicated. Examples of comets from the different treatment groups are shown. (C) Quantitation of the olive moment in > 200 cells from each treatment group is shown; error bars represent SEM. (D) Quantification of ROS levels in U2OS cells with endogenous ATM depletion and induced expression of WT, CL, or RX alleles of ATM, as measured by CellROX in triplicate. (E) Alkaline comet assays as in (B) showing effects of 1 day DRB treatment (20 μM). (F) Alkaline comet assays as in (B) with ATM depletion by shRNA and inducible expression of SETX. (G) DRIP-qPCR assays were performed in triplicate with primers specific for the BTBD19, β-actin, or EGR1 loci in U2OS cells depleted for endogenous ATM and expressing WT, CL, or RX alleles as indicated. Immunoprecipitations were performed in the absence of S9.6 antibody or with RNaseH treatment in vitro to verify antibody specificity. Levels of product were normalized to the level obtained in WT-expressing cells. Error bars indicate standard deviation. (H) DRIP-qPCR assays were performed in triplicate in human cells as in (G) at the BTBD19 and β-actin loci with the addition of 1mM NAC. (I) DRIP-qPCR assays were performed in triplicate in human cells as in (G) at the BTBD19 and β-actin loci with inducible expression of SETX as indicated. *, **, ***, and **** indicate p<0.05, 0.005, and 0.0005 by student t-test; NS = not significant.

Previous work has indicated that transcription may be involved in non-canonical forms of ATM activation (Bhatia et al., 2018; Britton et al., 2014; Sakasai et al., 2010; Sordet et al., 2009; Tresini et al., 2015) so we also tested the adenosine analog 5,6-dichloro-1-β-D-ribofuranosyl benzimidazole (DRB), an inhibitor of transcription elongation (Yankulov et al., 1995), and found that DRB treatment eliminates the single-strand breaks observed in the absence of ATM or with CL or RX allele expression (Fig. 3E).

We then considered the possibility that transcription-dependent single-strand breaks in ATM-deficient cells may involve R-loops, three-stranded RNA-DNA hybrid structures formed during transcription when an RNA transcript re-anneals to the template strand of the DNA (Crossley et al., 2019; Paull, 2019) since RNA-DNA hybrids have been shown to lead to strand breaks and genomic stability in many organisms (Aguilera and Gómez-González, 2017). To test for effects of R-loop removal in cells, we used inducible expression of the C-terminal half of the RNA-DNA helicase Senataxin (SETX), an enzyme that removes RNA species from genomic DNA and also promotes transcription termination (Lavin et al., 2013). We have previously found that recombinant SETX expression reduces high levels of R-loops in CtIP-deficient human cells and in *Δsae2* budding yeast (Makharashvili et al., 2018). Here we found that doxycycline-induced SETX expression drastically reduced the levels of single-strand breaks in comet assays with ATM depletion (Fig. 3F), consistent with the idea that R-loops are required for the DNA damage we observe in this assay.

To test this hypothesis further, we measured RNA-DNA hybrids at several genomic loci in human U2OS cells using DNA-RNA hybrid immunoprecipitation (DRIP) followed by quantitative PCR. A representative set of loci previously shown to be prone to R-loop formation (Bhatia et al., 2014; Hatchi et al., 2015; Makharashvili et al., 2018) was used for this analysis. These results showed an elevation of R-loops at the BTB19, β-actin, and EGR1 genes that was apparent with expression of the CL or RX alleles compared to expression of WT ATM (Fig. 3G). In contrast, expression of the R2579A/R2580A (2RA) allele, deficient in MRN activation (Lee et al., 2018), did not show significant differences from WT cells (Fig. S4). Appropriate controls were performed to ensure that the S9.6 antibody signal provides an accurate representation of R-loops under these conditions, including no-antibody samples and also incubation of the isolated genomic DNA with recombinant RNaseH before immunoprecipitation to test for RNA-DNA hybrid specificity. Preincubation of cells with NAC reduced the levels of R-loops observed at the BTB19 and β-actin loci (Fig. 3H), similar to the effects of NAC on single-strand breaks, indicating that ROS contributes to these lesions. Doxycycline-induced expression of SETX also reduced the levels of RNA-DNA hybrids observed in the absence of ATM, as expected (Fig. 3I).

To further investigate the relationship between transcription-dependent DNA damage and protein homeostasis in ATM-depleted cells, we tested for the effects of DRB treatment in the protein aggregation assay as in Fig. 1. Both CK2β and PSMB2 showed significantly higher enrichment in the aggregate fraction with ATM depletion in U2OS cells, which was reduced by DRB treatment (Fig. 4A, quantification of 3 replicates in Fig. 4B). We induced SETX expression in these cells and also saw a significant reduction in the aggregate level (Fig. 4C, D), consistent with the effects of SETX on single-strand breaks and R-loops. We also monitored ROS levels in these cells and found that SETX did not reduce the elevated ROS induced by ATM depletion (Fig. 4E), thus we infer that the R-loops and transcription-induced damage in these cells are downstream of ROS.

**Figure 4.**
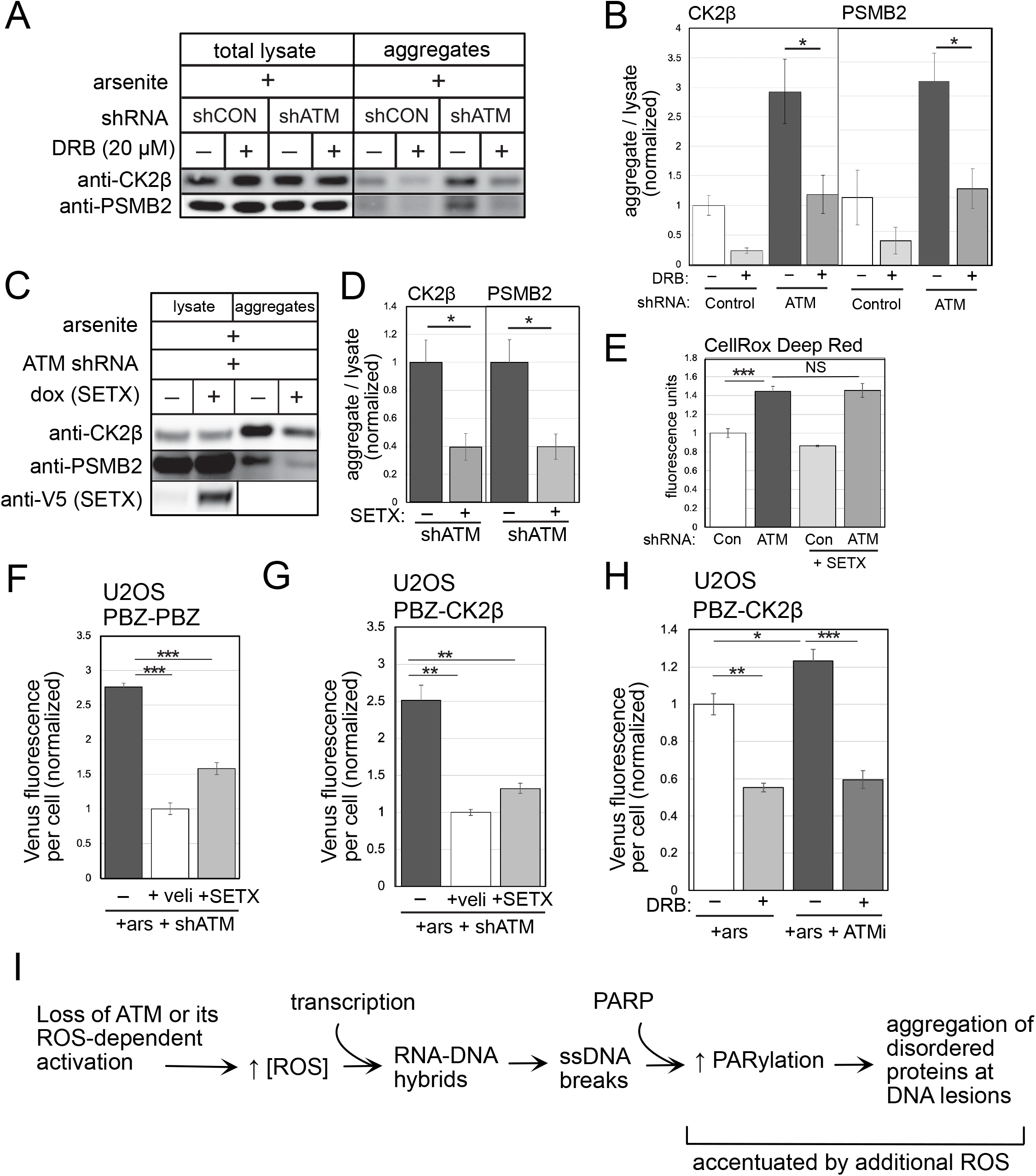
Active transcription and PARylation promote protein aggregation in human cells lacking ATM. (A) Detergent-resistant aggregates were isolated in U2OS cells shRNA-mediated depletion of ATM, arsenite (25 μM), and DRB (20 μM) added as indicated. Western blots for CK2β and PSMB2 in lysate and aggregate fractions are shown. (B) Three replicates of the experiment shown in (A) were performed and quantified. Levels of CK2β and PSMB2 in aggregate fractions normalized by lysate levels were quantified from each experiment and shown here relative to control cells. (C) Aggregation assays as in (A) with shRNA-mediated depletion of ATM, arsenite, and induction of SETX as indicated. (D) Three replicates of the experiment shown in (C) were performed and quantified. Levels of CK2β and PSMB2 in aggregate fractions normalized by lysate levels were quantified from each experiment and shown here relative to cells without SETX induction. Error bars indicate standard deviation. (E) Quantification of ROS levels in U2OS cells with ATM depletion and with SETX induction, as measured by CellROX in triplicate. (F) Fluorescence yield from the PBZ-PBZ PAR sensor measured in triplicate as in Fig. 2B with U2OS cells depleted for endogenous ATM, exposed to arsenite, and induced for SETX expression as indicated. (G) Fluorescence yield from the PBZ-CK2β sensor measured in triplicate as in Fig. 2E with U2OS cells depleted for endogenous ATM, exposed to arsenite, and treated with either veliparib or induced for SETX expression as indicated. (H) Fluorescence yield from the PBZ-CK2β PAR sensor as in Fig. 2E with U2OS cells exposed to arsenite, ATM inhibitor (ATMi, 1 μM AZD1390), and DRB as indicated. *, **, ***, and **** indicate p<0.05, 0.005, and 0.0005 by student t test; NS = not significant. (I) Diagram of events occurring in ATM-deficient or ATM oxidation activation-deficient cells. An increase in ROS, together with active transcription, generates R-loops and single-strand DNA breaks that hyperactivate PARP, leading to accumulation of disorder-prone proteins at DNA lesions.

Lastly, we measured PARylation using the live-cell sensors and found that the increase in PBZ-PBZ association caused by ATM loss was significantly reduced by SETX expression (Fig. 4F) or by RNaseH expression (Fig. S3), similar to the extent of reduction by the PARP inhibitor veliparib. The level of PBZ-CK2β association was also dramatic reduced by both SETX and by DRB (Fig. 4G, H). Thus, the elevated levels of PARylation observed in ATM-deficient cells occur in a ROS- and transcription-dependent manner under these conditions.

Taken together, this evidence shows that ATM-deficient and ATM oxidation-deficient human cells exhibit a number of pathological indicators of nuclear DNA damage that include high levels of single-strand breaks and elevated R-loops. Our evidence suggests that hyper-PARylation occurs as a result of these DNA lesions, and that direct association between oxidation-prone proteins and PAR polymer is caused by this series of events, ultimately generating the aggregation of disordered proteins at the sites of DNA lesions (Fig. 4I).

### An Mre11 ATLD mutant generates single-strand breaks and protein aggregation

Our model suggests that high ROS due to loss of ATM activation by oxidation is essential for the aggregation of the proteins we have monitored and also for neurodegeneration, considering that the R3047X mutation which specifically inhibits oxidative activation (Guo et al., 2010) is the causative genetic lesion in a subset of patients with A-T (Chessa et al., 1992; Gilad et al., 1998). One complication with this hypothesis, however, is the fact that rare mutations in Mre11 also generate A-T-Like Disorder (ATLD) which includes progressive cerebellar neurodegeneration (Stewart et al., 1999; Taylor et al., 2004). If protein aggregation and neurodegeneration are in fact independent of MRN-mediated activation of ATM, then the ATLD phenotype is inconsistent with this view.

To investigate this issue, we reconstituted an ATLD mutant allele in human U2OS cells (Fig. 5A). We depleted Mre11 with shRNA and expressed either shRNA-resistant WT Mre11 or a mutant allele containing both of the mutations present in ATLD3/4 patients, R572X and N117S (Stewart et al., 1999; Taylor et al., 2004), referred to here as “ATLD“. This combination of mutations results in expression of a truncated protein lacking the C-terminal 136 residues and is present at reduced levels compared to the WT allele. We confirmed that the replacement of endogenous Mre11 with the ATLD mutant compromised the activation of ATM by DNA double-strand breaks, as measured by camptothecin-induced Kap1 phosphorylation on S824 (Fig. 5A).

**Figure 5.**
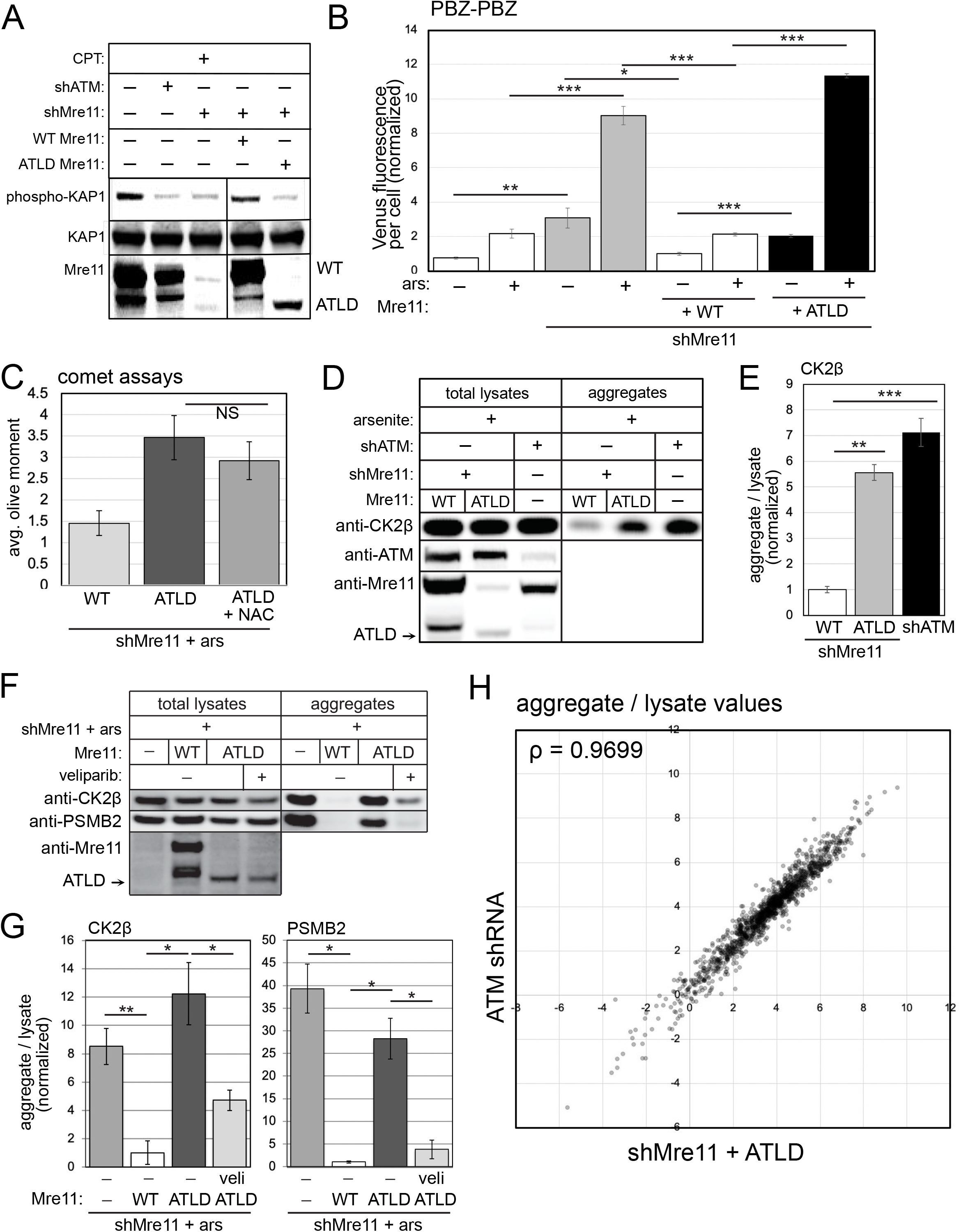
ATLD cells show elevated PARylation, single-strand DNA breaks and protein aggregation similar to ATM oxidation-deficient cells. (A) Endogenous Mre11 was depleted in human U2OS cells and WT or ATLD3/4 Mre11 was expressed, as indicated. The cells were exposed to camptothecin (10 μM for 1 hour) and blotted for phospho-S824 Kap1 as well as Mre11. (B) Fluorescence yield from the PBZ-PBZ PAR sensor was measured in triplicate as in Fig. 2B in U2OS cells depleted for endogenous Mre11, exposed to arsenite (25 μM), and expressing WT or ATLD Mre11 as indicated. Error bars indicate standard deviation. (C) Alkaline comet assays as in Fig. 1 with Mre11 depletion by shRNA and expression of ATLD Mre11 with NAC added as indicated in the presence of arsenite. Quantitation of the olive moment in > 200 cells from each treatment group is shown; error bars represent SEM. (D) Aggregation assays as in Fig. 1 with shRNA-mediated depletion of Mre11 and either WT or ATLD Mre11 expression compared to shRNA depletion of ATM as indicated. Total lysates and aggregate fractions were analyzed by western blot for ATM, Mre11, and CK2β. (E) Three replicates of the experiment shown in (D) were performed and quantified. Levels of CK2β and PSMB2 in aggregate fractions normalized by lysate levels were quantified from each experiment and shown here relative to cells expressing WT Mre11. Error bars indicate standard deviation. (F) Aggregate fractions and total lysates from 3 replicates of the experiment in (D) were analyzed by mass spectrometry. Aggregate values (normalized by lysate values) for each protein were determined (the mean of the 3 experimental replicates) and plotted for Mre11-depleted cells with ATLD expression (X axis) versus ATM shRNA-treated cells (Y axis). The Spearman correlation coefficient was calculated from this comparison to be 0.9699. (G) Aggregation assays as in Fig. 1 with shRNA-mediated depletion of Mre11 and either WT or ATLD Mre11 expression, with veliparib (10 μM) as indicated. Total lysates and aggregate fractions were analyzed by western blot for Mre11, CK2β, and PSMB2. (H) Three replicates of the experiment shown in (G) were performed and quantified. Levels of CK2β and PSMB2 in aggregate fractions normalized by lysate levels were quantified from each experiment and shown here relative to cells expressing WT Mre11. Error bars indicate standard deviation. *, **, ***, and **** indicate p<0.05, 0.005, and 0.0005 by student t test; NS = not significant.

Based on our results above indicating that PARP activation is a focal point that links ATM deficiency with protein aggregation, we tested the cells expressing WT or mutant Mre11 alleles for PARylation using the PBZ-PBZ sensor. This data shows that depletion of Mre11 generates higher levels of PARylation both in the presence and absence of arsenite (Fig. 5B). Expression of shRNA-resistant WT Mre11 eliminated this increase, while expression of the ATLD3/4 allele generated significantly higher PAR levels, similar to or even higher than Mre11 depletion alone. Consistent with this result, measurement of single-strand breaks using an alkaline comet assay showed that ATLD expression in the absence of endogenous Mre11 also generates higher levels of DNA damage compared to cells with WT full-length Mre11 expression (Fig. 5C). Interestingly, unlike with ATM depletion, the effect of Mre11 mutant expression on DNA breaks is not significantly reduced by antioxidant treatment (NAC)(Fig. 5C). We also do not see any change in ROS levels with depletion or mutation of Mre11 (Fig. S5).

We then measured protein aggregation using the ATLD Mre11 reconstituted cell lines, quantifying levels of CK2β lysates and in the detergent-resistant aggregate fraction. Here we observed increased aggregation of CK2β, compared to cells expressing WT Mre11, and similar to levels observed in cells with ATM depletion (Fig. 5D, E). Unlike with ATM depletion or inhibition, however, the aggregation observed with Mre11 loss was not reduced by NAC (Fig. S5). Nevertheless, ATLD-associated aggregates are reduced by veliparib (Fig. 5F, G), indicating a role for PARylation in promoting protein aggregation in the absence of normal Mre11 function, similar to what we observed with loss of ATM.

Lastly, we analyzed the lysates and detergent-resistant aggregate fractions from Mre11-depleted cells reconstituted with wild-type or ATLD Mre11, as well as ATM-depleted cells using label-free quantitative mass spectrometry. We observed a total of 2364 proteins present in all samples and calculated the abundance of each protein present in the total lysate as well as aggregate fractions, controlling the false discovery rate at 0.05 (Benjamini and Hochberg, 1995). From this analysis we determined that 1704 proteins showed significantly different aggregate values in Mre11 shRNA + ATLD expressing cells compared to WT Mre11 expressing cells (considered to be the control samples in this experiment), while 1584 targets were found to be significantly enriched in the aggregate fraction of ATM shRNA-treated cells compared to controls (Table S1). The aggregate levels (normalized by lysate amounts) of proteins showing significant differences in ATLD versus ATM shRNA cells are shown in Fig. 5H, where it is evident from visible inspection as well as the high Spearman correlation coefficient (ρ=0.9699) that the aggregation propensity of nearly all of these proteins is extremely similar in the absence of ATM function compared to the absence of Mre11 function.

From this analysis we conclude that loss of Mre11 function generates widespread protein instability extremely similar in breadth and magnitude to that observed with ATM depletion, and that expression of the ATLD allele cannot complement this deficiency. This is surprising given our previous observation that the specific ablation of MRN activation of ATM (in the 2RA mutant) does not generate protein aggregates (Lee et al., 2018), R-loops, single-strand DNA breaks, or high PARylation. The only explanation for this is that the loss of ATM activation by MRN is not functionally equivalent to the loss of the MRN complex itself.

### Protein aggregation is widespread in A-T patient cerebellum tissue

Up to this point we have focused on protein aggregation in ATM-deficient cell lines in culture but here we asked if the increase in protein aggregation also occurs in human A-T patients. To test for this, we acquired fresh-frozen cerebellum tissue from 21 deceased A-T patients, with ages at collection ranging from 8 to 47 years (Table S3). For comparison we also acquired 21 control samples, with age, sex, and race matched to patients. The tissue from each individual was homogenized in dry ice prior to lysate preparation, and was sufficient to perform two replicate aggregation preparations (Fig. 6A).

**Figure 6.**
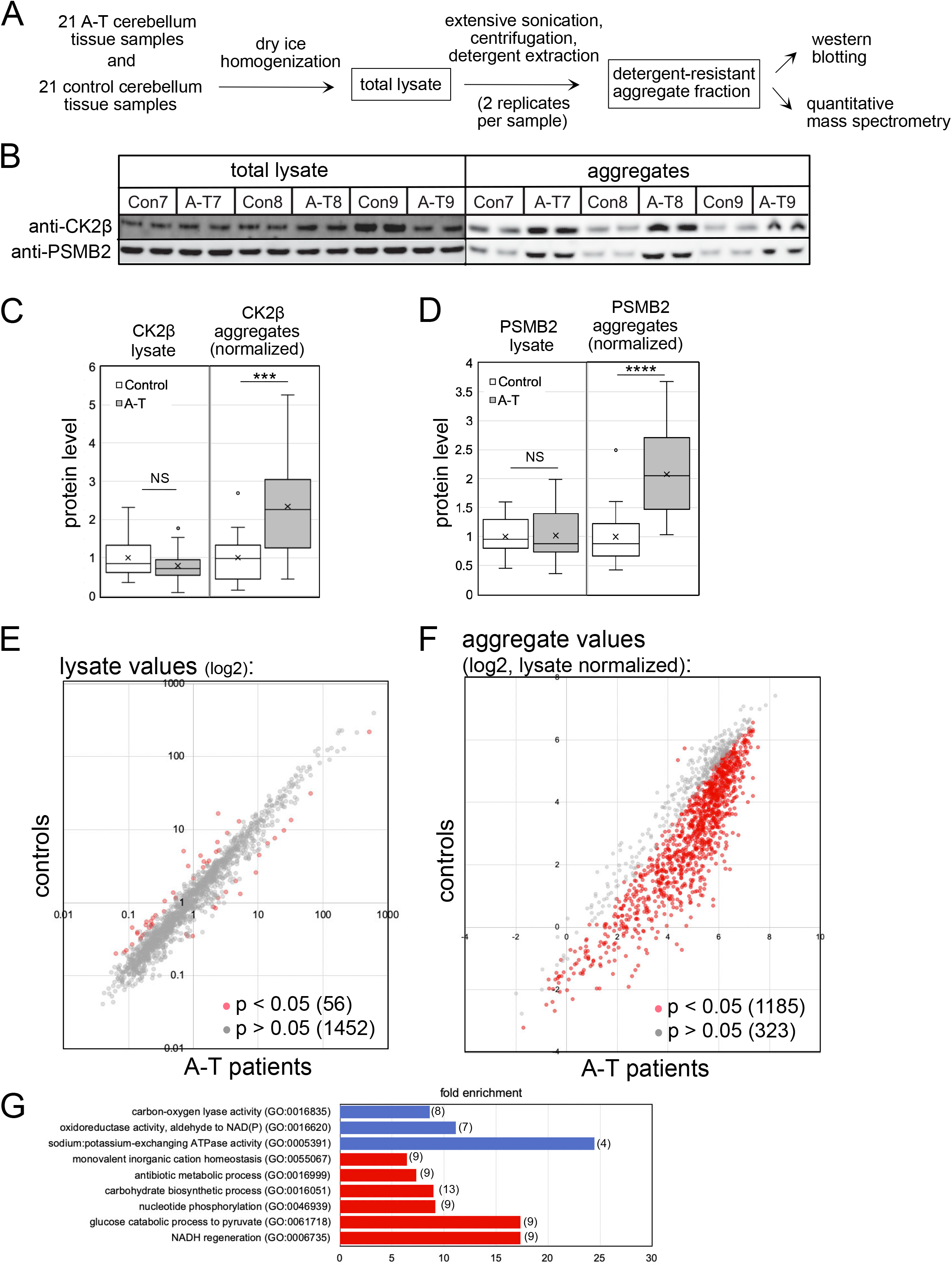
Global proteomics analysis of A-T patient cerebellum tissues and age-matched controls shows widespread protein aggregation. (A) Diagram of workflow: fresh-frozen cerebellum slices from 21 A-T patients ages 8 to 50 were obtained from the NIH neurobiobank, as well as age/gender/race-matched control samples. Each tissue was homogenized and filtered to generate a lysate and split into 2 replicates, each of which was further processed by sonication, centrifugation, and detergent extraction to generate an aggregate fraction, and all samples were analyzed by western blotting and mass spectrometry. (B) Representative western blot showing levels of CK2β and PSMB2 in lysates as well as aggregate fractions. (C, D) Quantification of western blotting results from all 42 lysates (replicates averaged) and 42 aggregate samples (normalized by lysate amounts) as indicated. All values are also normalized to levels in control group. (E). Levels of 1508 proteins found in the lysates of control and A-T patient samples as determined by mass spectrometry. Red shows proteins with significant differences between patient and control groups. (F) Aggregate levels from the 1508 proteins quantitated in A-T patients relative to controls, normalized by lysate. Red shows proteins with significant differences between patients and controls. (G) Gene Ontology terms and fold enrichment of proteins identified in (F), number of genes in parentheses. Error bars indicate standard deviation. *, **, ***, and **** indicate p<0.05, 0.005, and 0.0005 by student t test; NS = not significant.

The lysate and aggregate fractions from each of the 42 individuals (with technical replicates for each) were analyzed first by western blotting, as done previously with cell line-derived samples. An example of a portion of this dataset is shown in Fig. 6B, blotted for CK2β and PSMB2. The results show generally similar levels of the proteins in lysates, but enrichment of these factors in the detergent-resistant aggregate fractions. Analysis performed for all of the samples show a statistically significant increase (2 to 2.5-fold) in CK2β and PSMB2 aggregates when comparing all 21 A-T patients as a group to the control individuals, while no differences were observed in lysate levels (Fig. 6C, D).

We then analyzed all samples by quantitative mass spectrometry which identified a total of 4350 proteins, using a minimum of 2 unique peptides identified with high confidence as a threshold. Replicates of the lysate and aggregate fractions were analyzed, with a high level of similarity between replicates (average Spearman rho correlation coefficients for normalized aggregate values were 0.945 and 0.898, respectively, for controls and patients). 2460 proteins were identified in all lysate samples from all individuals, while 2577 proteins were identified in all aggregate samples. From this analysis we identified 1508 proteins for which we obtained a complete set of quantitative values for all lysate and all aggregate samples in all individuals in both replicates, which was used to calculate normalized aggregate levels for each protein (aggregate/lysate ratio)(Table S4).

The total lysates from patients and control individuals show relatively few statistically significant differences (56 proteins among the 1508 proteins with complete data for all individuals, Fig. 6E, Table S4). Nevertheless, it is notable that the proteins in this group include IP3R1, INPP5A, and CBLN1 (Fig. S6); loss of any of these factors individually in mammals causes spinocerebellar ataxia (Dudding et al., 2004; Huang et al., 2012; Klar et al., 2017; van de Leemput et al., 2007; Yang et al., 2015).

In contrast, the number of proteins with significantly different levels of aggregates in A-T patient cerebellum samples compared to controls was very high (1158 of 1508 total)(Fig. 6F, Table S4). A vast majority of these (1156 of 1158) exhibited higher aggregate levels in A-T patients compared to controls, reminiscent of the results we observed in human cell lines (Fig. 1).

We previously showed that aggregation-prone proteins observed in the absence of ATM oxidative activation in cultured human cells exhibit higher predicted disorder scores when measured by the TANGO algorithm for aggregation propensity (Fernandez-Escamilla et al., 2004; Lee et al., 2018). The aggregation scores for the 1156 proteins found to be more enriched in A-T brain samples were also analyzed in this manner and were found to have even higher scores (average of 2792.5 compared to the average for the total proteome, 1910.4) than the aggregates previously identified.

Analysis of the aggregate-prone proteins identified in the cerebellum samples shows enrichment for metabolic enzymes, including glycolysis-related enzymes such as GAPDH, pyruvate kinase PKM, triosephosphate isomerase, and alpha-enolase (top 10% of hits analyzed for gene ontology term enrichment shown in Fig. 6G). Other enriched categories include NADH regeneration and sodium/potassium transport proteins such as the ATP1A and ATP1B families and calcium transport and binding proteins, including ATP2B1/2, SLC8A1/2, and CALB1 (Fig. S6). Oxidoreductases are also generally enriched on this list, including pyruvate dehydrogenases (PDHA1 and PDHB), aldehyde dehydrogenases (ALDH2), and peroxireductase (PRDX1). Chaperones are present in the enriched fraction, most notably the HSP70 chaperones HSP70 and HSC70 and GRP78/HSPA5, all of which bind to and regulate the folding of misfolded proteins. Lastly, it is notable that proteins associated with other forms of neurodegeneration are found enriched in the aggregate list, including Ataxin-10, a protein required for the survival of cerebellar neurons and mutated in Spinocerebellar ataxia type 10 (Teive et al., 2011), and alpha-synuclein, well-known for its aggregation propensity in Parkinson’s Disease (De Mattos et al., 2020).

Comparison of the aggregation-prone polypeptides in U2OS cells (including proteins found to be aggregated both ATM-depleted as well as ATLD cells), U87-MG cells, and in A-T patient samples identifies a core set of 189 aggregation-prone proteins (Fig. S7). This comparison indicates similar global aggregation propensity in cultured cells and in patient cerebellum samples, particularly when considering the fact that the overlap in total proteome among this diverse set of cell lines and brain tissue includes only 705 polypeptides.

### Aggregate and PARylation levels distinguish A-T patients from controls

Analysis of the cerebellum tissue shows that the protein aggregation we observed in human cells in culture with depletion of ATM is also present at significant levels in A-T patient cerebellum tissue. We also used the normalized aggregate levels for all of the individuals to perform unsupervised hierarchical clustering (Fig. 7A). Consistent with the strong bias for aggregation observed in patients as a group (Fig. 6F), this analysis separates most of the patients from the controls. However, it is clear that there is variation among the patients, and a group of 5 patients (A-T 3, 6, 17, 19, and 21) cluster separately from the rest. Included in this subset are the two youngest patients (A-T 3 and 21), both 8 years old at the time of sampling.

**Figure 7.**
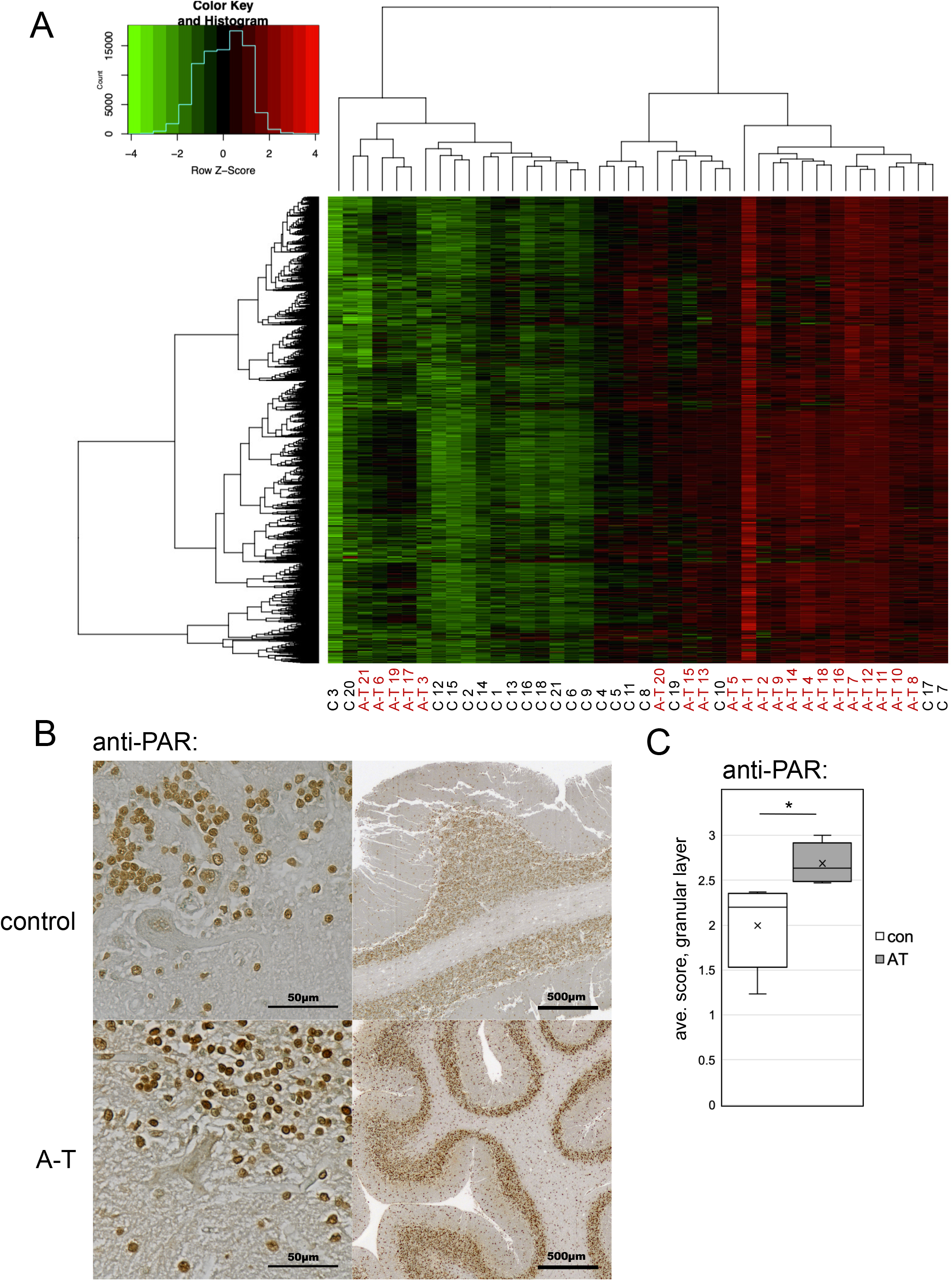
Hierarchical clustering of A-T patient and control protein aggregate values separates most patients from the control population. (A) Heat map of aggregate values from Figure 1 after unbiased hierarchical clustering. A-T patient and control identifiers are listed on the bottom. (B) Cerebellum tissue (formalin-fixed) from a control and an A-T patient (#6), analyzed by immunohistochemistry with an antibody directed against poly-ADP-ribose, with HRP-conjugated secondary antibody and 3,3’Diaminobenzidine, using methyl green as a counterstain, at two resolutions as indicated. (C) Subjective scoring of the level of PAR staining of cells in the granular layer of 5 A-T patient cerebellum samples (5, 6, 12, 18, 21) compared to age-matched controls was performed using a 0 to 3 scale of blinded samples by 3 individuals. Error bars indicate standard deviation. *, **, ***, and **** indicate p<0.05, 0.005, and 0.0005 by student t test; NS = not significant.

A subset of the patient cerebellum tissues was also available as formalin-fixed samples. Several of these were examined using immunohistochemistry (IHC) for overall organization of the cerebellum, as well as for levels of poly-ADP-ribose using an anti-PAR antibody (Fig. 7B). Analysis of patient samples showed that all contained identifiable molecular, granular, and Purkinje cell layers, thus the patient tissue is not grossly dissimilar to that of the controls with respect to cerebellum organization (A-T 6 shown here as an example). Blinded scoring of 5 patient tissue samples and 5 controls revealed a significant increase in overall PAR staining intensity in the patient group as compared to the controls (Fig. 7C).

## Discussion

### Widespread protein aggregation is a characteristic of A-T patient cerebellum tissue

All of the prevalent neurodegenerative disorders in humans are associated with a loss of protein homeostasis that ultimately results in aggregation of specific proteins in disease-specific patterns in the brain (Bourdenx et al., 2017). A-T has not previously been considered in this way, but here we present evidence showing that this cerebellum-specific neurodegenerative disorder also exhibits widespread protein aggregation, affecting more than 1000 polypeptides in human patient cerebellum tissue. Proteins found in these aggregates range from metabolic enzymes to signaling components that are, on average, much higher in predicted intrinsic disorder compared to the total proteome.

In other neurological diseases where mutations in specific disorder-prone factors predispose individuals to neurodegeneration, the presence of the mutant aggregating protein clearly plays a causal role in tissue dysfunction (Currais et al., 2017; Soto and Pritzkow, 2018). The progression toward disease endpoints is also influenced by the levels of protein dysfunction, natural aging, as well as levels of oxidative stress and DNA damage (Madabhushi et al., 2014; Maynard et al., 2015).

In A-T patient cerebellum tissues, protein aggregation is clearly elevated when comparing patients as a group to control individuals. The insoluble polypeptides identified at higher levels in patients have overall higher predicted levels of intrinsic disorder and are enriched for specific classes of enzymes. Hierarchical clustering of patients and controls by aggregate values groups most of the patients together, but five patients (including the two youngest) show a different pattern of modest aggregation levels (Fig. 7). It is possible that these patterns represent earlier stages of the disease, with relatively few aggregating proteins that are stochastically distributed. If so, these individuals illustrate patterns early in the degenerative process and could yield further insights into the initial molecular events that drive tissue degeneration.

### A-T cerebellum contains lower levels of proteins associated with dominant cerebellar ataxia syndromes

The total proteome quantifications we performed on the patient and control lysate samples did not show a large number of differences compared to the aggregate comparisons. However, we found several proteins with reduced levels in A-T patients overall that are very important in other forms of cerebellar ataxia. For instance, the intracellular calcium channel IP3R1 (expressed from the *ITPR1* gene) is 7-fold lower in A-T patients relative to the control distribution (Fig. S6).

*ITPR1* deletions in humans cause a dominant form of spinocerebellar ataxia, SCA15/16 (Dudding et al., 2004; van de Leemput et al., 2007), and missense mutations in these genes are linked to congenital nonprogressive ataxia known as SCA29 (Huang et al., 2012; Klar et al., 2017). IP3R1 is regulated by levels of inositol 1, 4, 5-triphosphate, a signaling molecule hydrolyzed by the INPP5A phosphatase. Deletion of the *INPP5A* gene causes cerebellar degeneration in mice (Yang et al., 2015), emphasizing the important role for this signaling pathway in Purkinje cells. We also found that INPP5A levels are low in A-T patients (2.8-fold decreased relative to the control mean). Lastly, carbonic anhydrase 8 (CA8), a binding partner of IP3R1, is also significantly decreased in A-T patient cerebellum tissue. Missense mutations in *ITPR1* that abolish IP3R1 binding to CA8 are pathological mutations in SCA29 (Ando et al., 2018). Mutant ataxin-2 and ataxin-3 proteins responsible for the more common SCA2 and SCA3 disorders, respectively, have also been shown to interact abnormally with IP3R1 (Kasumu and Bezprozvanny, 2012). Taken together, the observations in our dataset strongly suggest the possibility of deficient inositol phosphate-regulated calcium signaling in A-T patients. It is notable in this regard that the early A-T literature documented “aberrant sensing” of calcium by A-T cells in culture, reduced duration of calcium currents in A-T neuronal cells, as well as deficiencies in phosphoinositol metabolism (Famulski and Paterson, 1999; Famulski et al., 2003; Khanna et al., 1997; Yorek et al., 1999), but the molecular basis of these observations has yet to be resolved.

### PARP activity promotes protein aggregation in the absence of ATM oxidative activation

Our results show that poly-ADP-ribosylation by PARP1 and 2 is important for the aggregation of proteins observed in the absence of ATM function. PARP has previously been shown to be hyperactivated in ATM-deficient cells (Fang et al., 2016), although the depletion of NAD^+^ caused by PARP overactivation was implicated as the critical factor in that study. We have not observed any effects of NAD^+^ replenishment in our work (using nicotinamide riboside as an NAD^+^ donor), thus we conclude that it is the hyper-PARylation itself and not the depletion of NAD^+^ that generates the protein aggregates in ATM-deficient cells.

PARylation at sites of DNA damage has already been shown to attract intrinsically disordered proteins (Altmeyer et al., 2015; Chen et al., 2018; Hong et al., 2013; Mastrocola et al., 2013; Rulten et al., 2014; Singatulina et al., 2019). In a few specific cases, phosphorylation of these disordered proteins by ATM has been shown to disperse them from PARylation sites (Chen et al., 2018; Gardiner et al., 2008), suggesting that ATM function is important for reversal of PAR-induced insoluble protein domains. We propose that loss of ATM combined with excessive PARylation and disordered protein accumulation leads ultimately to the irreversible loss of protein solubility that we observe in the tissues.

PARP has also been shown to be specifically neurotoxic in specific disease states. PARP1 hyperactivity promotes cerebellar ataxia in mice in the absence of XRCC1 for instance (Hoch et al., 2017), also a situation in which excess single-strand breaks generate PARP overactivation. PARP also promotes the neurotoxicity of several RNA-binding proteins in a Drosophila model of ALS (Duan et al., 2019), and has been shown to accelerate alpha-synuclein-dependent neurodegeneration in Parkinson’s disease (Kam et al., 2018). All of this evidence is consistent with a role for PAR in generating stable, toxic protein aggregates. PARP activity does facilitate single-strand break repair, however (Gupte et al., 2017), and deletion of ATM and PARP1 simultaneously in the mouse generates synthetic lethality (Murcia et al., 2001). It is possible that PARP activity (or even hyperactivity) is required for the rapid cell divisions during early mammalian embryogenesis in the context of ATM-deficiency, but then becomes toxic in later stages of growth by promoting the assembly of protein aggregates.

### Transcription-associated lesions in ATM-deficient cells generate single-strand breaks, PARP hyperactivation, and protein insolubility

Our results strongly suggest a role for ATM in preventing global single-strand breaks, with important components of this phenotype contributed by excess ROS as well as active transcription. The canonical view of ATM activation via DNA damage requires DNA double-strand breaks, an activation pathway we have previously demonstrated in vitro with purified components and many groups have confirmed in mammalian cells (Lee and Paull, 2005; Paull, 2015; Shiloh, 2003). Others have suggested more recently that ATM can also be activated by single-strand breaks in DNA where it acts to delay S phase entry (Khoronenkova and Dianov, 2015), and that base excision repair (BER) efficiency is decreased upon loss of ATM protein, coincident with the accumulation of oxidized protein and increases in nuclear proteasome activity (Poletto et al., 2017). Changes in BER-mediated repair of single-strand breaks in the absence of ATM may contribute to the sponataneous breaks we have observed.

In addition, ATM activation can be induced by Top1 poisons such as camptothecin, which was reported to generate ATM activation in a transcription-dependent manner (Sakasai et al., 2010; Sordet et al., 2009), although double-strand breaks were invoked in this scenario as the activating lesion. Based on this evidence as well as observations of Top1 and Top2 conjugates accumulating in the absence of ATM function (Alagoz et al., 2013; Katyal et al., 2014; Yamamoto et al., 2016) or Mre11 (Hoa et al., 2016), we overexpressed Tdp1 or Tdp2 in ATM-depleted cells since these enzymes directly promote resolution of topoisomerase conjugates (Pommier et al., 2014). However, this did not alleviate the protein aggregation we observed (data not shown). We note that Top2a was slightly elevated in aggregate fractions from ATLD-expressing U2OS cells compared to WT Mre11-expressing cells but was unchanged in ATM-depleted cells (Table S1). Top1 levels were not statistically different in either case, nor was it found to be elevated in aggregate fractions from A-T patients (Table S4). Lastly, camptothecin treatment does not induce aggregation in the absence of ATM function (data not shown), despite the fact that Top1 conjugates are strongly induced under these conditions (Pommier, 2006). We conclude that Top1 conjugates are not likely to be the critical lesion that induces the aggregation.

The fact that elevated ROS is required for hyperactivation of PAR, accumulation of single-strand breaks, and aggregation in the absence of ATM oxidative activation suggests an alternative model where ROS-induced transcriptional stalling generates R-loops and, ultimately, single-strand breaks (Fig. 4I). This has been suggested in other contexts where active transcription in the presence of oxygen stress was shown to generate R-loops that trigger transcription-coupled DNA repair (Teng et al., 2018). Our data showing that SETX overexpression alleviates not only the R-loops but also single-strand breaks, hyper-PARylation, and protein aggregation shows that RNA-DNA hybrids are a critical intermediate in this pathway. Consistent with this model, mutations in SETX are linked to neurodegenerative disorders AOA2 (ataxia with oculomotor apraxia type 2) and amyotrophic lateral sclerosis type 4 (ALS4), and expansions in C9ORF72 that are causative for ALS are also known to generate pathological R-loops (Wang et al., 2015).

It is not clear how loss of ATM leads to R-loop accumulation. ATM phosphorylation of splicing-related targets was shown to alter irradiation-induced spliceosome dynamics (Tresini et al., 2015), which could be important considering the close relationship between splicing efficiency and R-loop formation (Santos-Pereira and Aguilera, 2015). R-loops were not found to be present at higher levels in ATM-deficient mice (Yeo et al., 2014), although the mouse models of A-T do not show overt cerebellar degeneration or ataxia (Barlow et al., 1996; Elson et al., 1996; Xu et al., 1996).

### ATLD is not functionally equivalent to loss of ATM activation by MRN

The association between the MRN complex and ATM was initiated by the seminal discovery of *MRE11* mutations in patients diagnosed with A-T, giving rise to the “A-T-like Disorder” designation (Stewart et al., 1999). Subsequent work showed that MRN is required for ATM activation by double-strand breaks in mammalian cells and in purified systems (Lee and Paull, 2004, 2005; Uziel et al., 2003), solidifying the model that the clinical features observed in ATLD are attributable to a loss of ATM activation via MRN. It is certainly true that MRN plays a critical role in facilitating ATM activation at double-strand breaks; however, a series of recent observations suggest that the neurodegeneration in ATLD patients may not be related to this specific function.

First, three cerebellar ataxia patients were identified in France with previously unreported biallelic mutations in the MRE11 gene, diagnosed with ATLD (Fiévet et al., 2019). Unlike other ATLD patients, cells from these individuals showed no defects in ATM activation despite relatively low levels of MRN complex. This data indicates that the ATLD phenotype does not necessarily include a global loss of ATM-mediated DNA damage responses.

Second, we identified a separation-of-function allele of ATM (2RA) that is specifically deficient in ATM activation via the MRN complex and DNA double-strand breaks in vitro with purified components as well as in human cells (Lee et al., 2018). Expression of the 2RA allele of ATM in place of the WT allele did not generate high ROS or protein aggregation, even in the presence of additional oxidative stress, while alleles that block the oxidative pathway show high levels of aggregates. Considering the correlation between protein aggregates and A-T status shown in this study, the results with the 2RA allele suggests that loss of MRN-stimulated ATM activity is not the critical event leading to loss of protein homeostasis observed in ATM-deficient cells.

Third, the analysis we performed in this study suggests that loss of the MRN complex, accentuated by expression of ATLD alleles (a combination of ATLD3/4 R572X and N117S used in this work), generates many of the same functional consequences as loss of ATM, including high single-strand breaks, hyper-activation of PARP, and protein aggregation. A detailed comparison of the levels of each misfolded polypeptide shows remarkable convergence between ATLD and loss of ATM, analogous to the neurological similarities between ATLD and A-T patients. Combined with the additional observation that the ROS activation-specific allele of ATM (R3047X) also causes neurodegeneration (Chessa et al., 1992; Gilad et al., 1998), this suggests that ATLD shares a defect with ATM-deficient cells and cells lacking ATM oxidative activation.

Interestingly, cells lacking Mre11 (either with or without ATLD allele expression) show the same loss of protein homeostasis observed with ATM-deficient cells, but ROS does not appear to play a critical role. For instance, the antioxidant NAC does not significantly reduce the levels of single-strand DNA breaks in ATLD cells, nor does it block protein aggregation. ROS levels are also not elevated in Mre11-depleted cells, in contrast to ATM-depleted cells. We do not know the origin of single-strand DNA breaks in cells lacking MRN, but this will be an important area of future study.

Overall, our investigation of human cerebellum tissue as well as cell lines is consistent with the proposal that loss of protein homeostasis is a marked feature of both ATM and Mre11 deficiency in human cells. Results in tissue culture clearly show that the activities of PARP enzymes are important for this phenomenon, stimulated by single-strand breaks that are created in a transcription-dependent manner. More detailed analysis of PARylation targets and dynamics will help us to understand the trajectory of events from initial lesion to the ultimate loss of cerebellum function that occurs in this untreatable disease.

## Acknowledgements

We thank members of the Paull laboratory for helpful discussion and help with blinded scoring, as well as Domenico Delia for critical suggestions, to Vishwanath Iyer for the glioblastoma cell line, and to Rajashree Deshpande and Oshadi Wimalarathne for important reagents and imaging contributions. We are indebted to the families of the A-T patients and control individuals who contributed the autopsy material for research use. We acknowledge the Howard Hughes Medical Institute for funding this project and Cancer Prevention and Research Institute grant RP170628.

## Author Contributions

J-H. L., S.W.R., and N.E. conducted experiments, evaluated data, and contributed to the editing of the manuscript. T.T.P generated reagents, evaluated data, and wrote the manuscript.

## STAR Methods

### Gene expression constructs

Recombinant ATM expression in U2OS cells was performed with pcDNA5-FRT/TO-intron derived vectors containing shRNA-resistant ATM alleles, including WT (pTP3540), C2991L (pTP3362), R2579A/R2580A (pTP4011), D2889A (pTP4012), and R3047X (pTP3552). Depletion of endogenous ATM was performed by incubating cells with lentivirus containing shRNA toward ATM (sc-29761-SH, Santa Cruz Biotechnology). The C-terminus of Senataxin was expressed from a pcDNA5-FRT-TO derivative containing C-term-V5 SETX, a.a. 1851 to 2677 (pTP3531)(Makharashvili et al., 2018). RNaseH was expressed from a pcDNA5-FRT-TO derivative containing *E. coli* rnhA fused to mCherry. The original rnhA plasmid was a gift from Patrick Calsou (Addgene #60365)(Britton et al., 2014). The PBZ-PBZ sensor was derived from pBiFC-PBZ-VC and pBiFC-PBZ-VN (Addgene #110648 and 110646, respectively) but combined into one bacmam plasmid, pTP4623, using the backbone of pAceBac1 (Bieniossek et al., 2008). The original pBiFC PBZ constructs were gifts from Chris Lord (Krastev et al., 2018). The PBZ domain fused to VC in the PBZ-PBZ plasmid pTP4623 was also replaced with human CK2β, creating the PBZ-CK2β sensor in pTP4658. These plasmids were used to generate bacmids using standard methods with the bac-to-bac system (Life Technologies), and virus was produced in Sf21 insect cells. Depletion of endogenous Mre11 was performed with lentivirus expressing shRNA (5′-ACAGGAGAAGAGAUCAACUUUG-3′) from the H1 promoter using the backbone from lenticrispr v2 (Addgene #52961), a gift from Feng Zhang (Sanjana et al., 2014), with Cas9 deleted, pTP4359. WT shRNA-resistant Mre11 with a C-terminal Flag tag was expressed from a pcDNA5-FRT-TO derivative, pTP4099. A mutant version of this plasmid containing Mre11 N117S and R572X mutations was generated using quikchange mutagenesis (Agilent) to create the ATLD expression plasmid pTP4225. Cloning details and sequence files are available upon request.

### Cell culture and recombinant protein induction

U2OS T-Rex FLP-in cells containing plasmids with ATM and MRE11 genes were cultured in Dulbecco’s Modified Eagle Medium (DMEM, Invitrogen) supplemented with 10% fetal bovine serum (FBS, Invitrogen) containing 15 μg/ml Blasticidin (A1113903, Life Technology), 100 units/ml penicillin-streptomycin (15140-122, Life Technology), and 200 μg/ml Hygromycin (400052-50ml, Life Technology). Depletion of endogenous ATM was performed by incubating cells with lentivirus containing shRNA cassettes overnight and selecting with media containing 1 μg/ml puromycin (Invitrogen) for 5-7 days. To induce genes introduced through the FLP-in system, doxycycline (1 μg/ml; Thermo Fisher, #BP-2653-5) was added to the medium as final concentration 3 days before treatment with arsenite (25 mM; Sigma, #S7400-500G). Lentivirus was prepared in HEK-293T cells as previously described (Lee et al., 2018). U87-MG glioblastoma cells were obtained from Vishwanath Iyer and were grown under the same conditions as U2OS cells.

### Split venus PBZ sensors

U2OS cells were plated in T75 flasks until cells reached 70% confluency. 5 ml baculovirus second amplification supernatant containing either PARylation sensor or PBZ/CK2ß sensor bacmam was added to each flask with 5ml of DMEM containing 10% FBS. A day after infection, cells were plated in multiple wells of a 6-well plate at approximately X% confluence. Cells were treated with either 25μM arsenite, 1μM AZD1390, 1mM NAC, 1μg/ml Doxocycline or 10μM Veliparib as indicated in the figure legends. The following day, cells were harvested, washed with cold PBS containing 0.9mM Ca^2+^, 0.5mM Mg^2+^, and resuspended in Triton X-100 extraction buffer (0.5% Triton X-100, 20 mM Hepes-KOH (pH 7.9), 50 mM NaCl, 3 mM MgCl2, 300 mM Sucrose) for 2 minutes. Extracted samples were recovered by centrifugation at 1200Xg at 4°C for 3 minutes, then resuspended in 500 uL cold PBS and kept on ice until FACS analysis (no more than 2 hours). Flow Cytometry: Extracted cells were analyzed with a BD LSRII Fortessa Flow cytometer using fluorescence detection parameters for Alexa Fluor^®^ 488 (excitation 488nm; emission 519nm) for detection of split venus. Extracted cells were gated according to forward scatter and side scatter in order to remove any partial or non-extracted cells. For all experiments at least 10,000 events measured. Average and standard deviation were generated with the median of fluorescence value from three biological replicates.

### Comet assay

U2OS cells were grown in DMEM (10% FBS) media in the presence of doxycycline (1 mg/ml) in 6-well plates for 3 days and treated with NAC (1 mM) as indicated in the figure legends for 18 hours before harvesting. Alkaline comet assays were performed using OxiSelect Comet assay Kit (Cellbiolabs, #STA-350) following the manufacturer’s protocol. Samples were observed under a Zeiss Axiovert 200M fluorescence microscope and analyzed with Fiji imageJ program.

### ROS measurements

Cells treated with CellRox deep red reagent (Thermo Fisher, #C10422) at a final concentration of 5 mM for 30 min at 37°C. Cells were harvested with trypsin, washed with PBS, and then transferred to 5 ml of polystyrene round bottom tubes (VWR, #60818-496) in 1 ml of PBS for analysis by flow cytometry according to the manufacturer’s protocol.

### DNA-RNA immunoprecipitation (DRIP) assay

Cells were harvested with trypsin, resuspended in 5 ml of PBS supplemented with 0.5% SDS and 1 mg/ul Proteinase K (Gold biotechnology, #P-480-500) as a final concentration, and digested at 37°C overnight. Genomic DNA was isolated and performed DRIP assay as previously described (Makharashvili et al., 2018).

### Aggregation assay

To prepare cell lysates, the pellets were resuspended in lysis buffer (20 mM Na-phosphate pH 6.8, 10 mM DTT, 1mM EDTA, 0.1% Tween 20, 1 mM PMSF, and EDTA-free protease inhibitor mini tablets (Thermo Fisher, #A32955)) and rotated at 4°C for 30 min. Cells were sonicated in a 4°C water bath-based sonicator (Bioruptor: 8 times at level 4.5 and 50% duty cycle) and centrifuged for 20 min at 200g at 4°C. The concentration of protein in the supernatant was calculated using the Bradford reagent (Thermo Fisher, #23236) and adjusted to a same concentration for all samples. Protein aggregates were pelleted at 16,000g for 20 min at 4°C. After removing supernatants, protein aggregates were washed 2 times with NP-40 buffer (20 mM Na-phosphate pH 6.8, 2% NP-40, 1mM PMSF, and EDTA-free protease inhibitor mini tablets), sonicated 6 times at level 4.5 and 50% duty cycle, and centrifuged at 16,000g for 20 min at 4°C. Aggregated proteins were washed in wash buffer (20 mM Na-phosphate pH 6.8, 1mM PMSF, and EDTA-free protease inhibitor mini tablets), sonicated 4 times at level 3 and 50% duty cycle, and boiled in 2X SDS sample buffer. Resuspended aggregated proteins were analyzed by either western blotting or mass spectrometer.

### Cerebellum tissue homogenization and aggregate preparation

Fresh-frozen cerebellum tissue (vermis, approximately 1 g) was obtained for 21 A-T patients ranging in age from 8 to 50 years as well as age/gender-matched controls. Tissue samples were pulverized with 15 ml powdered dry ice for 2 min in a commercial blender on the highest setting. The tissue and dry ice mixtures were brushed into disposable weigh dishes then poured into 50 ml conical tubes and placed on ice prior to cell lysis. All steps were performed in a 4°C cold room with materials kept chilled on dry ice. Powdered brain tissues were incubated with lysis buffer (see Aggregation assay, above) for 20 min on ice, sonicated 5 times at level 4.5 and 50% duty cycle, and centrifuged for 5 min at 200g. Supernatants were procced to the standard aggregation assay by starting with sonication 8 times at level 4.5 and 50% duty cycle.

### Filter aided sample preparation

Frozen tissue lysates and pellets were kept in SDS loading buffer prior to experiment. Sample preparation was performed as previously described (Lee et al., 2018)with following change. 8μg of protein for lysates or whole samples for pellets were diluted to 800μL of UA (8M Urea, 0.1M Tris-HCl pH8.8) total volume. After filter aided sample preparation, label free quantification LC-MS/MS was performed by the proteomics facility in the University of Texas at Austin following previously described procedures. The raw data was processed by Proteome Discoverer 2.2.

### Statistics testing and analysis

Results from Proteome Discoverer 2.2 were further refined by removing common known contaminates and any proteins identified with less than two unique polypeptides. Then refined data was normalized by western blot results of corresponding samples using CK2β amount in order to correct for variation in sample loading. Pellet over lysate values were generated by dividing pellet values with corresponding lysate values then taking log (base 2) of the results. Proteins with missing data were dropped for the analysis. For U2OS cells, P-values comparing control to shATM or shMre11 were generated using Welch’s T-test. Two technical replicates from brain samples were averaged before the Welch’s T test comparing control patients to A-T patients. Benjamini-Hochberg procedure was used in order to control for multiple hypothesis testing using 0.05 FDR. Heatmap and hierarchical clustering were generated using R studio version 1.2.5001. with heatmap.2 function under gplots package in conjugation with colorRampPalette function in RColorBrewer package. The hierarchical clustering was performed using default hclust method with “euclidean” distance calculation and “complete” clustering.

### Immunohistochemistry

Formalin-fixed cerebellar tissue blocks were dehydrated, paraffinized, and cut into 4 μm thick sections using standard methods. Tissue sections were mounted on charged glass slides, deparaffinized with xylene then rehydrated with graded alcohols. Antigen retrieval was performed by incubation with Proteinase K for 20 min at 37°C. After rinsing in TBST, sections were incubated overnight at 4°C with primary antibody for PAR (Abcam PN: ab14459, 1:100 in IHC World diluent PN: IW1000). Endogenous peroxidase activity was quenched with 3% H_2_O_2_ in water for 15 min before incubation with poly-HRP secondary antibody (Thermo Fisher PN: 32230, 1:400 in TBS-1% BSA) for 1 hr at room temperature. Sections were stained using BD DAB substrate kit (PN: 550880), counterstained with methyl green, dehydrated in ethanol, cleared in xylene and mounted with DPX mountant (Millipore Sigma). A negative control for non-specific secondary antibody binding was generated following the above procedure omitting primary antibody addition. Nuclear staining with DAB was observed in slides incubated with PAR antibody while only methyl green counterstain was present in negative control. Five images of each A-T patient or control tissue at 40X magnification were scored by 3 individuals, with patient/control identity blinded. The scoring was performed as previously described (Emre et al., 2020), with a scale of 0 (not detected), 0.5 (very low), 1 (low), 1.5 (low to moderate), 2 (moderate), 2.5 (moderate to strong), and 3 (strong), analyzing only cells in the granular layer.

### Data Availability

Mass spectrometry raw data or original files for other experiments are available upon request.

