## Supplemental materials for "Poly-ADP-ribosylation drives loss of protein homeostasis in ATM and Mre11 deficiency"

**Supplementary Figures 1 - 7**

**Supplementary Table 1**: Quantitation of polypeptides in U2OS cells with ATM shRNA depletion or Mre11 shRNA depletion with either wild-type Mre11 or ATLD expression by mass spectrometry. Lysates and aggregate fractions shown separately.

**Supplementary Table 2:** Quantitation of polypeptides in U87-MG cells with ATM shRNA depletion, ATM shRNA depletion plus NAC treatment, or control cells, by mass spectrometry. Lysates and aggregate fractions shown separately.

**Supplementary Table 3**: List of A-T patients and controls.

**Supplementary Table 4**: Quantitation of polypeptides in human cerebellum samples by mass spectrometry. Lysates and aggregate fractions shown separately.

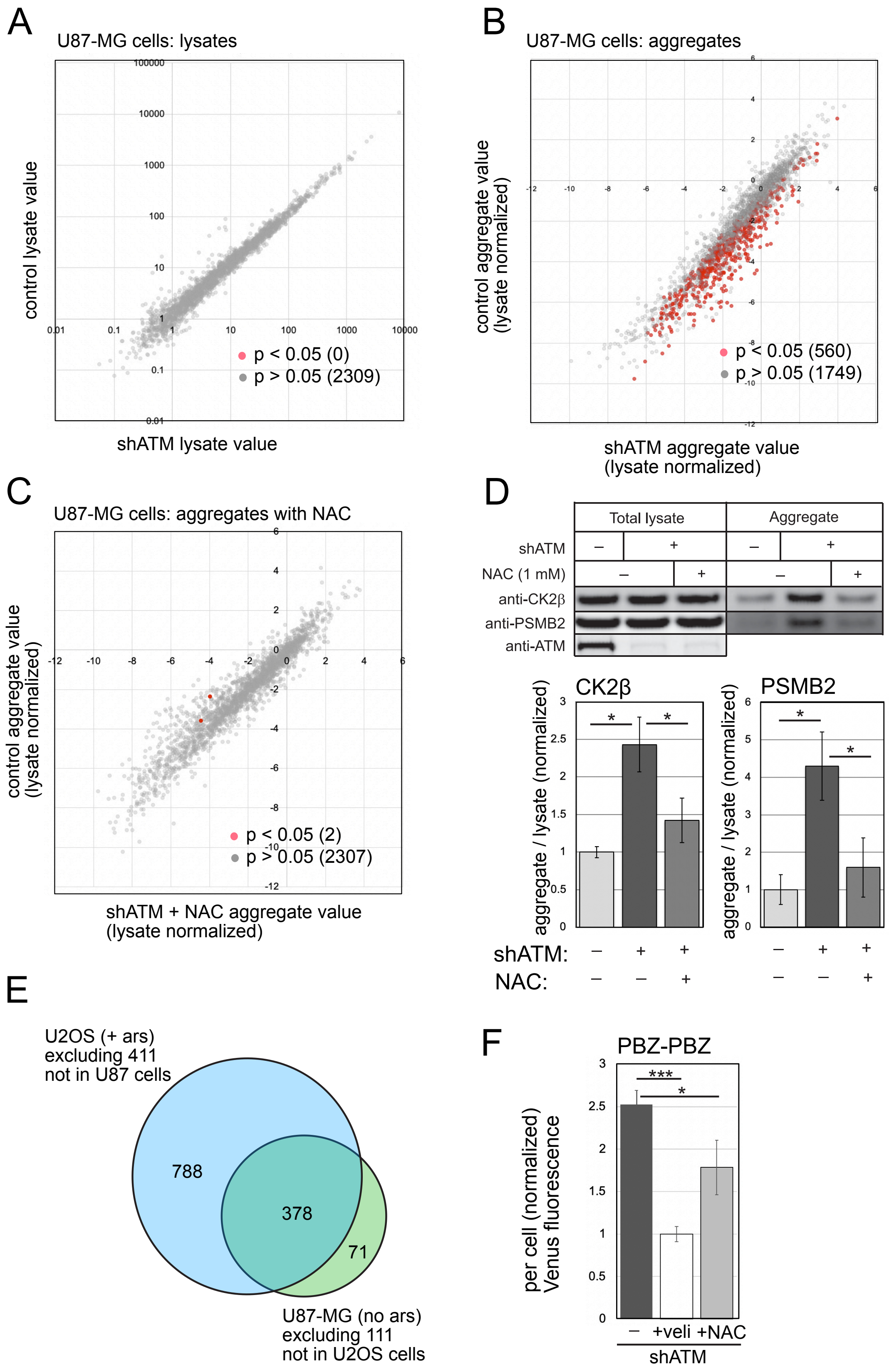

A
U87-MG cells: lysates
control lysate value
shATM lysate value
p < 0.05 (0)
p > 0.05 (2309)
B
U87-MG cells: aggregates
control aggregate value
(lysate normalized)
shATM aggregate value
(lysate normalized)
p < 0.05 (560)
p > 0.05 (1749)
C
U87-MG cells: aggregates with NAC
control aggregate value
(lysate normalized)
shATM + NAC aggregate value
(lysate normalized)
p < 0.05 (2)
p > 0.05 (2307)
D
Total lysate
Aggregate
shATM
NAC (1 mM)
anti-CK2β
anti-PSMB2
anti-ATM
CK2β
PSMB2
aggregate / lysate (normalized)
shATM:
NAC:
E
U2OS (+ ars)
excluding 411
not in U87 cells
788
378
71
U87-MG (no ars)
excluding 111
not in U2OS cells
F
PBZ-PBZ
per cell (normalized)
Venus fluorescence
+veli
+NAC
shATM

**Supplementary Figure 1. Loss of ATM in human glioblastoma cells (U87-MG) results in hyper-PARylation and protein aggregation.** (A) Human U87-MG glioblastoma cells were depleted of ATM by shRNA and compared to control cells (3 biological replicates for each). Cell lysates were prepared and analyzed by mass spectrometry, identifying 2309 polypeptides in all samples. The levels of each protein in control versus ATM-depleted cells are shown, with non-significant differences in grey (2309) and significant differences in red (0), after FDR control at 0.05. (B) Detergent-resistant aggregates were prepared from control and ATM-depleted cells described in (A), and characterized by mass spectrometry. The levels of each protein in control versus ATM -depleted cells are shown, with non-significant differences in grey (1749) and significant differences in red (560), after FDR control at 0.05. (C) Detergent-resistant aggregates were prepared and characterized from control and ATM-depleted cells as in (B) but ATM-depleted cells were pre-incubated with NAC (1 mM). The levels of each protein in control versus ATM-depleted cells are shown, with non-significant differences in grey (2307) and significant differences in red (2), after FDR control at 0.05. (D) Cell lysates and aggregates from (C) were analyzed by western blot, with antibodies specific for CK2β, PSMB2, and ATM as indicated. Bottom panels: Three replicates of the experiment were performed and quantified. Levels of CK2β or PSMB2 in aggregate fractions normalized by lysate levels were quantified from each experiment and shown here relative to control cells. (E) Overlap between aggregated proteins identified in U87-MG with those identified in U2OS cells with arsenite treatment, excluding proteins not present in both types of cells. (F) FACS results from three replicates showing the mean fluorescence yield per cell from U87-MG cells expressing the PBZ-PBZ PAR sensor with ATM shRNA depletion and either veliparib (10 μM) or NAC (1 mM) treatment as indicated. At least 10,000 cells were measured in each replicate. Fluorescence yield was normalized to veliparib-treated cells.

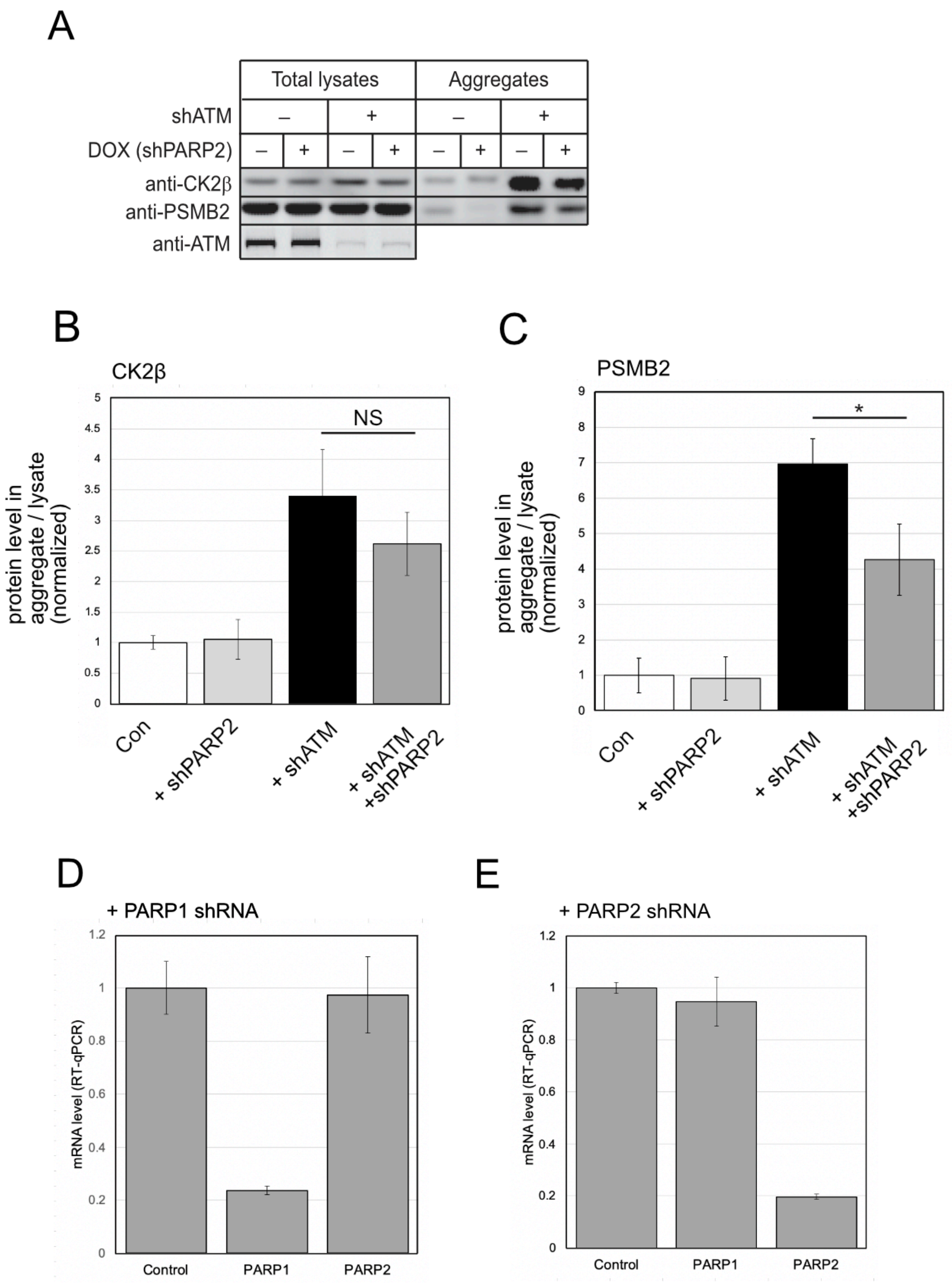


**Figure S2. Reduction of PARP2 partially lowers aggregate levels in ATM-depleted cells.** (A) U2OS cells were depleted of endogenous ATM as in Fig. 1 and doxycycline-induced shRNA specific for PARP2 was expressed. Aggregation assays with cells depleted of ATM and concurrently depleted for PARP2 were performed. Levels of CK2β and PSMB2 in aggregate fractions normalized by lysate levels were quantified from each experiment and shown here relative to control cells. (B, C) Three replicates of the experiment shown in (A) were performed and quantified. Levels of CK2β and PSMB2 in aggregate fractions normalized by lysate levels were quantified from each experiment and shown here relative to cells without ATM depletion. Error bars indicate standard deviation. (D, E) Levels of PARP1 and PARP2 mRNA in cells lines used in (A-C) as determined by RT-qPCR, normalized to levels in control cells. * indicates $p<0.05$ by student t test; NS = not significant.

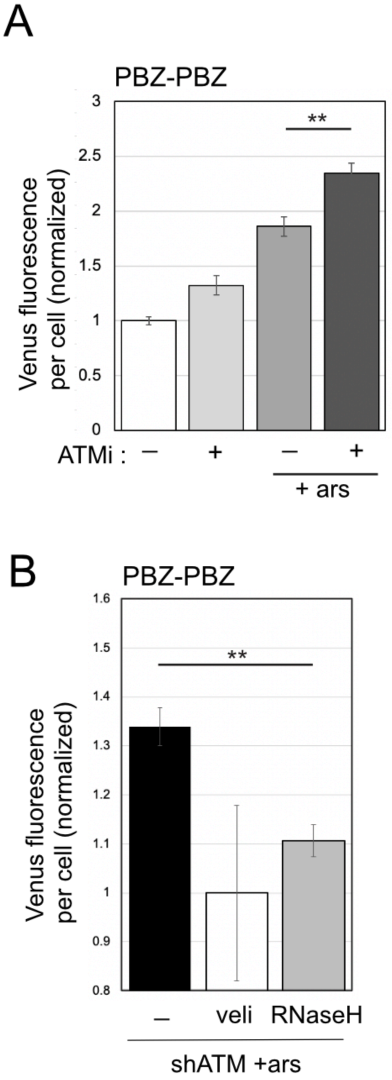


**Supplementary Figure 3. Increased PARylation with ATM depletion does not depend on active cell replication and can be reduced by overexpression of RNaseH.** (A) FACS results from three replicates showing the mean fluorescence yield per cell from cells expressing the PBZ-PBZ PAR sensor but incubated in media lacking serum for 7 days. $G_1$ DNA content of cells was verified by propidium iodide staining (data not shown). ATM inhibitor (ATMi, 1 µM AZD1390) and arsenite (25 µM) were added to cells one day before measurement of at least 10,000 cells per replicate. Fluorescence yield was normalized to control cells. (B) FACS results from three replicates showing the mean fluorescence yield per cell from cells expressing the PBZ-PBZ PAR sensor with ATM depletion and arsenite (25 µM) as indicated, with veliparib treatment (10 µM) or overexpression of bacterial RNaseH as indicated, with levels normalized to veliparib. At least 10,000 cells were measured in each replicate. Fluorescence yield was normalized to control cells. *, **, ***, and **** indicate $p<0.05$, 0.005, and 0.0005 by student t test; NS = not significant.

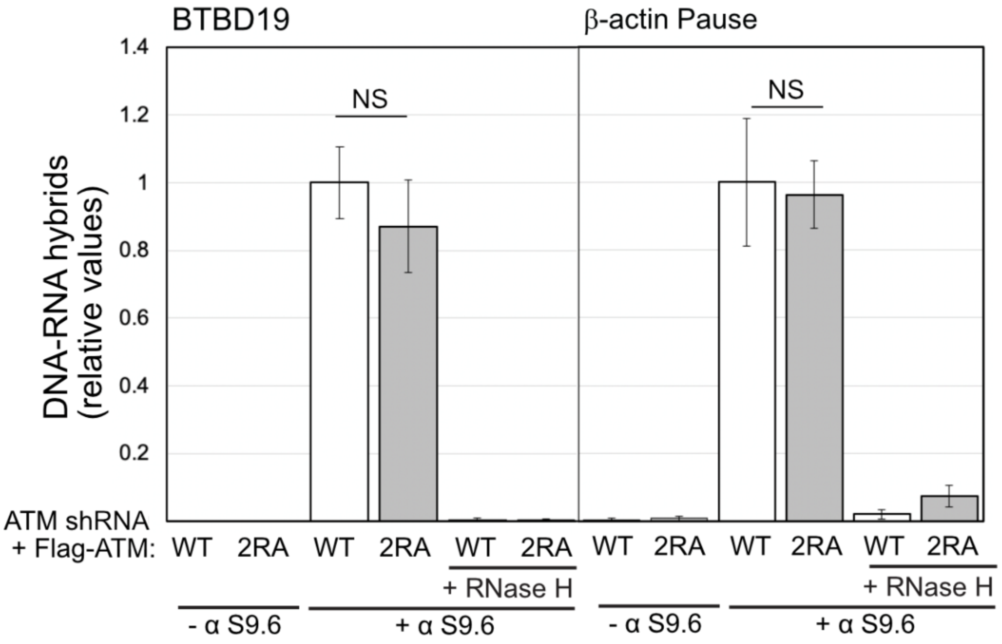


**Supplementary Figure 4. Loss of MRN activation of ATM does not increase R-loops in human cells.** U2OS cells were depleted of endogenous ATM as in Fig. 1 and shRNA-resistant alleles of wild-type (WT) or R2579A/R2580A (2RA) alleles of ATM were inducibly expressed from a genomic locus as indicated. DRIP-qPCR assays were performed in triplicate with primers specific for the BTBD19 or β-actin loci. Control reactions were also performed in the absence of S9.6 antibody or with RNaseH treatment in vitro to verify antibody specificity. Levels of product were normalized to the level obtained in WT-expressing cells. Error bars indicate standard deviation. NS = not significant.

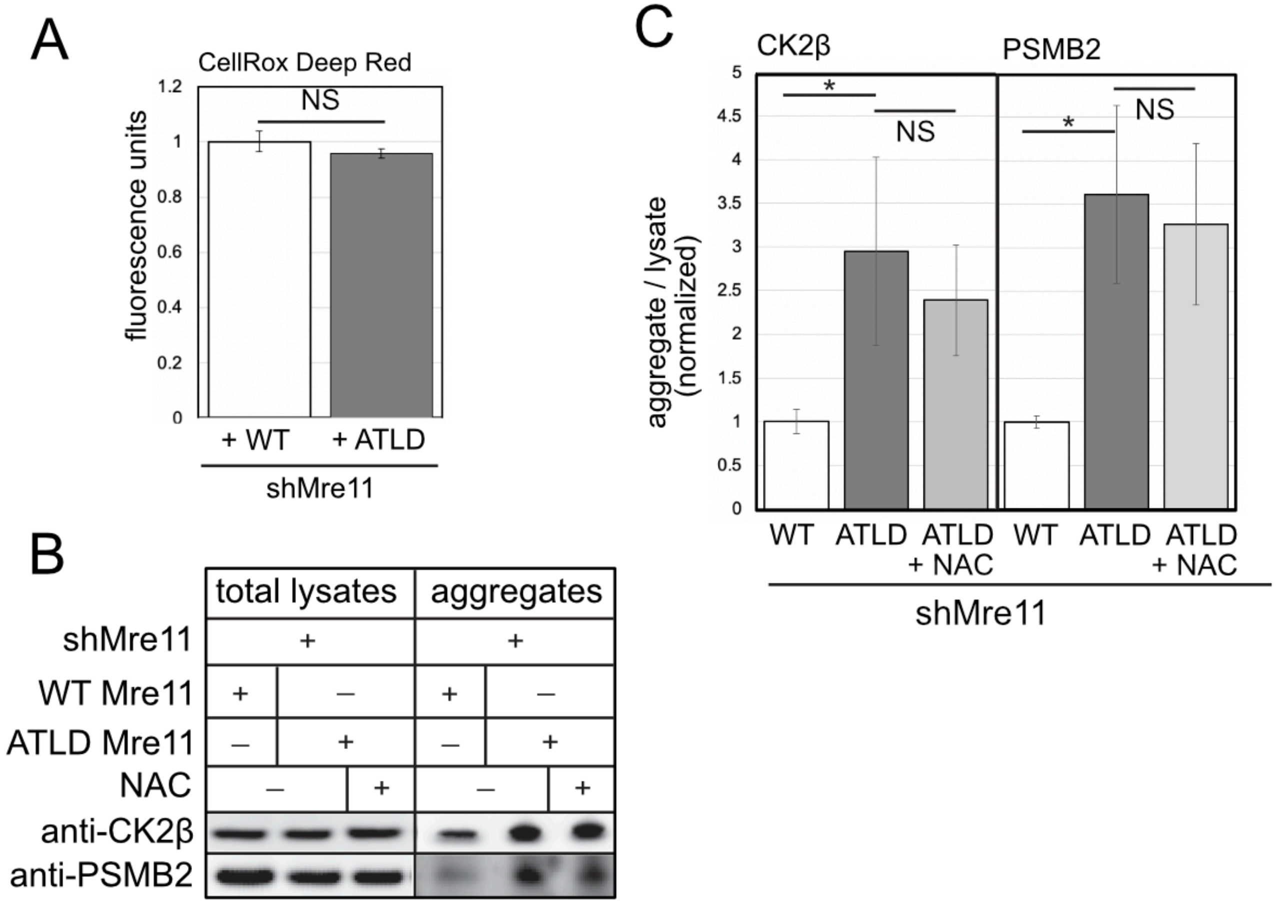


**Supplementary Figure 5. Expression of ATLD Mre11 does not increase ROS, and reduction of ROS does not lower aggregate levels in ATLD-expressing cells.** (A) Total ROS was measured in U2OS cells depleted of endogenous Mre11 and overexpressing either wild-type (WT) or ATLD Mre11 alleles, measured in triplicate using CellROX deep red, with levels normalized to WT-expressing cells. (B) Aggregation assays as in Fig. 1 with shRNA-mediated depletion of Mre11 and either WT or ATLD Mre11 expression compared to shRNA depletion of ATM and addition of NAC (1 mM) as indicated. Total lysates and aggregate fractions were analyzed by western blot for CK2β and PSMB2. (C) Three replicates of the experiment shown in (B) were performed and quantified. Levels of CK2β and PSMB2 in aggregate fractions normalized by lysate levels were quantified from each experiment and shown here relative to cells expressing WT Mre11. Error bars indicate standard deviation. *, **, ***, and **** indicate $p<0.05$, 0.005, and 0.0005 by student t test; NS = not significant.

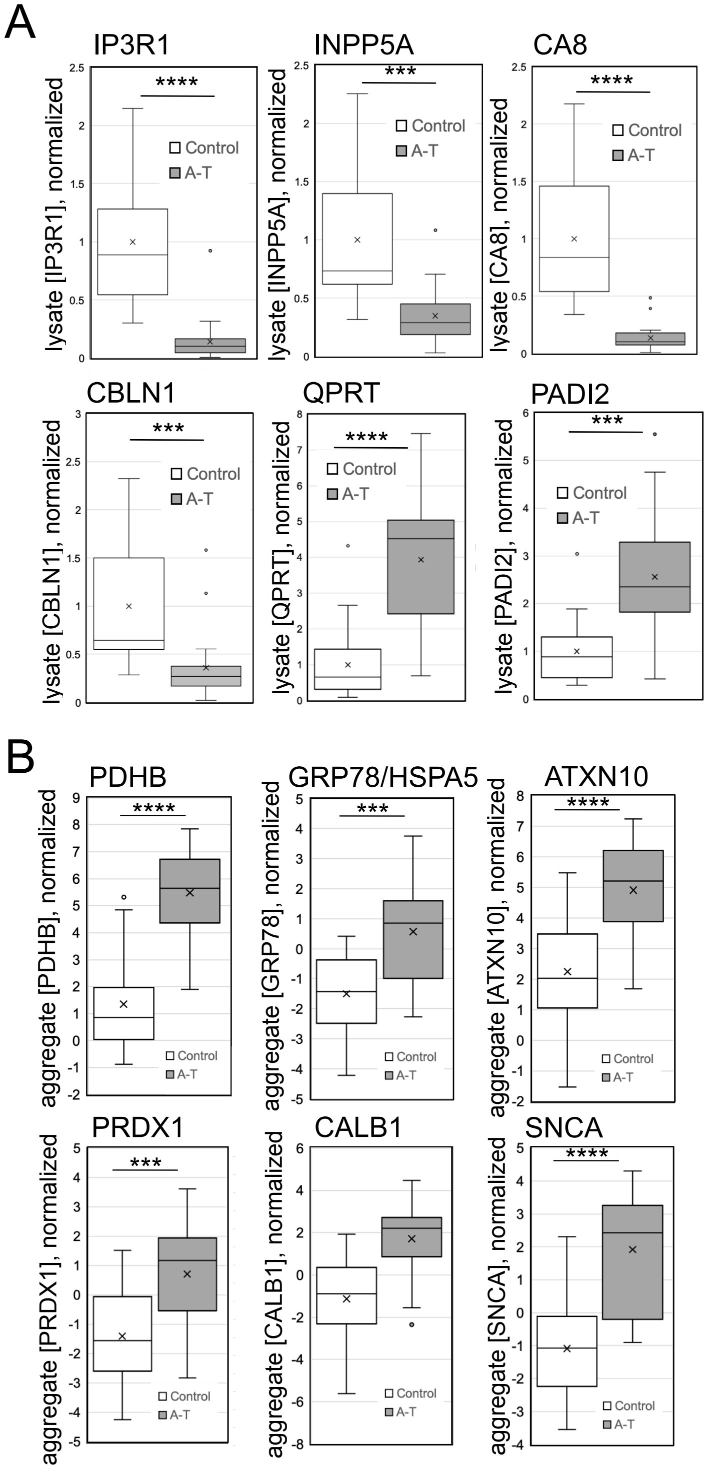


**Supplementary Figure 6. Examples of individual protein level differences in A-T and control cerebellum tissues.** (A) Lysate values for proteins with significant differences between A-T patients and controls, with control average normalized to 1. Error bars indicate standard deviation. (B) Aggregate values for proteins with significant differences between A-T patients and controls. Aggregate values are normalized by lysate amount for each individual. Error bars indicate standard deviation. *, **, ***, and **** indicate p<0.05, 0.005, and 0.0005 by student t-test; NS = not significant.

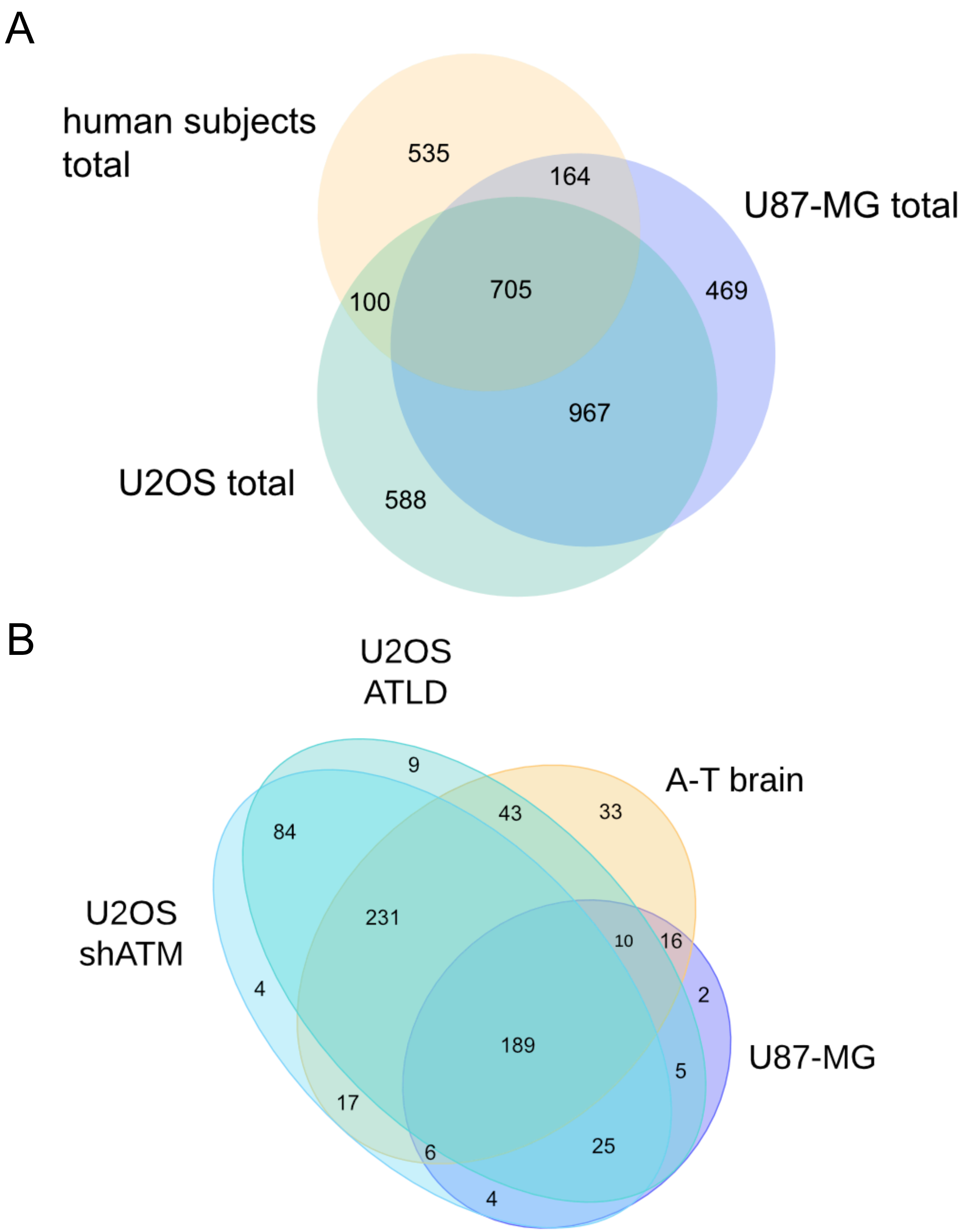


**Supplementary Figure 7. A core set of aggregation-prone proteins in human cells.** (A) Overlap between total proteomes identified by mass spectometry in U2OS, U87-MG cells, and human cerebellum samples. "Total" here refers only to proteins identified in both lysate and aggregate fractions, in all samples, across all replicates. (B) Overlap between proteins with significant differences in pellet/lysate ratio in ATLD (shMre11 + ATLD), shATM groups in U2OS cells, U87-MG glioma with shATM, and aggregation-prone factors identified in A-T patient cerebellum samples. Welch t test corrected for FDR 0.05 with Benjamini-Hochberg. Only proteins present in all biological systems are shown. Samples from U2OS were collected with arsenite exposure.
